# Inferring time of infection from field data using dynamic models of antibody decay

**DOI:** 10.1101/2022.10.03.510698

**Authors:** Benny Borremans, Riley O Mummah, Angela H Guglielmino, Renee L Galloway, Niel Hens, K C Prager, James O Lloyd-Smith

**Author notes:** **Preprint**, This manuscript is a preprint and has not been peer-reviewed. It is currently under review at a peer-reviewed scientific journal. We welcome any feedback, through the comments section on the bioRxiv website or by emailing the corresponding author.

## Abstract

Studies of infectious disease ecology often rely heavily on knowing when individuals were infected, but estimating this time of infection can be challenging, especially in wildlife. Time of infection can be estimated from various types of data, with antibody level data being one of the most promising sources of information. The use of antibody levels to back-calculate infection time requires the development of a host-pathogen system-specific model of antibody dynamics, and a leading challenge in such quantitative serology approaches is how to model antibody dynamics in the absence of experimental infection data. Here, we present a way to do this in a Bayesian framework that facilitates the incorporation of all available information about potential infection times. We apply the model to estimate infection times of Channel Island foxes infected with *Leptospira interrogans*, leading to reductions of 51-92% in the window of possible infection times. Using simulated data, we show that the approach works well across a broad range of parameter settings and can lead to major improvements of infection time estimates that depend on system characteristics such as antibody decay rate and variation in peak antibody levels after exposure. The method substantially simplifies the challenge of modeling antibody dynamics in the absence of individuals with known infection times, opens up new opportunities in wildlife disease ecology, and can even be applied to cross-sectional data once the model is trained.

## Introduction

Knowing when individuals got infected with a pathogen can dramatically boost insights into infectious disease biology, at both population and within-host scales (Handel and Rohani, 2015; Pepin et al., 2017). This knowledge allows estimation of incidence (the number of new infections over time) and force of infection (the rate at which susceptible individuals become infected), quantities that are fundamental to understanding and modeling transmission dynamics (Heisey et al., 2006; Held et al., 2019; Hens et al., 2010) and developing mitigation strategies (Caley and Hone, 2004; Weitz et al., 2020).

Knowledge of individual infection times is also relevant to a wide range of pathogen-related factors, including interpretation of the time course of clinical signs of disease (Hawley et al., 2011), vaccine efficacy (Antia et al., 2018), risk factors for infection (Borremans et al., 2011; Pepin et al., 2019), pathogen spillover (Smith et al., 2014), effects of disease on wildlife health and survival (Tersago et al., 2012), host immunity (Epstein et al., 2013) and tracing infection sources (Craft, 2015). However, even though a variety of data sources can theoretically be used to estimate infection time (e.g. clinical signs of disease, antibody concentration, outbreak seasonality, contact tracing), there are significant challenges that limit the widespread adoption of time-of-infection approaches, particularly in wildlife. Key challenges for estimating individual time of infection include how to incorporate individual variation in response to infection (Simonsen et al., 2009; Teunis et al., 2002), how to integrate different data sources (Borremans et al., 2016), how to deal with interval-censored data (Wilber et al., 2020), how to model the anamnestic response to reinfection (Pothin et al., 2016) and how to deal with antibody cross-reactivity. A currently unresolved major challenge is how to model biomarker dynamics when there is no population of individuals with a known infection time, such as a group of animals infected experimentally and tracked longitudinally; this challenge is particularly common in wildlife studies.

Models of the dynamics of biomarkers such as antibodies or pathogen DNA/RNA constitute the foundation of most time-of-infection estimation methods (Brookmeyer and Gail, 1988; Gilbert et al., 2013; Teunis et al., 2016), and are central to the rapidly expanding field of quantitative serology (Boni et al., 2019; Pepin et al., 2017; Teunis et al., 2012). The presence and concentration of such biomarkers can contain information about whether and when an individual has been infected (Borremans et al., 2016), the degree of immunity (Röltgen et al., 2021), infection severity (Vaughn et al., 2000), and whether and for how long they are infectious (Hardestam et al., 2008; Prager et al., 2020). Crucially, a biomarker can be used for such purposes only after its relevant properties have been quantified and when a model exists of how its presence or concentration correlates with the information of interest (e.g. time since infection). For example, a model of immunity to reinfection with rabies virus in vaccinated wildlife suggests that protective immunity should occur when the level of specific neutralizing antibodies is above a certain threshold (Moore et al., 2017). While biomarker models can range from purely conceptual to applied data-driven mathematical frameworks, they must exist before interpretation of new data is possible.

Antibody dynamics can be a particularly rich source of information about time of infection. Following infection, the humoral immune response results in the production of different types of antibodies that are produced at different rates and in different quantities. Antibody levels decline after reaching a peak level shortly after infection, and this decay follows a particular functional form, with a characteristic rate. When this functional form and rate are known, antibody levels measured at some later point can potentially be used to determine how long ago an individual was infected (Boni et al., 2019; Teunis et al., 2012). This in turn opens up the possibility to improve difficult-to-collect data on incidence in the population (Pepin et al., 2017; Wilber et al., 2020) and perhaps to estimate whether and for how much longer an individual is immune to reinfection (Borremans et al., 2015).

Models of antibody dynamics typically represent a general pattern in antibody levels: an increasing phase (often ignored because it is short), a peak level, and a decay phase (Teunis et al., 2016). Describing this pattern quantitatively requires specification of a functional form and associated parameters, and there are specific data requirements for estimating these parameters. The optimal situation for parameter estimation is the availability of experimental data where the time of infection is known for multiple individuals, combined with frequent longitudinal sampling of each individual until antibodies are no longer detectable. For example, experimental infection of the African rodent *Mastomys natalensis* with an arenavirus, followed by frequent sampling for the entire lifetime, enabled the development of an antibody dynamic model that could then be used to estimate time of infection of wild rodents based on a limited number of samples (Borremans et al., 2016, 2015). Similarly, experimental data on influenza A in snow geese and mallards have been used to model the antibody response following infection, and subsequently estimate infection times and population-level force of infection (Pepin et al., 2017).

However, in most studies of wildlife diseases, such experimental infection data are not feasible to collect, and infection times in the field are unknown. This is particularly problematic because periods between sampling can be long, and sampling sizes are typically small (Gilbert et al., 2013). This has been a major reason that quantitative serology methods have not yet been widely adopted in wildlife disease ecology (Gilbert et al., 2013). A standard approach to determining an animal’s time of infection is to take the midpoint between the interval bounded by the most recent time at which an individual is known to be antibody-negative and the first positive sample, or to consider this interval as a uniform probability distribution for infection (i.e. any time is equally possible). For example, a study on cowpox virus in field voles (*Microtus agrestis*) assumed a uniform infection probability over a period from 2 weeks prior to the last negative result until 2 weeks prior to the first positive result, based on the assumption (i.e. a basic model) that antibodies are detectable 2 weeks after infection and remain detectable for longer than the period between sampling events (Begon et al., 2009). Similarly, the time of seroconversion to Rift Valley fever virus in livestock used the midpoint between negative and positive samples taken at 1- to 2-month intervals (van den Bergh et al., 2019), which were subsequently used to estimate incidence over time. Intervals of multiple months or even years are common in wildlife systems, leading to potentially large errors surrounding time of infection estimates, especially when a uniform distribution between sampling times is assumed. This offers opportunities for improvement, as quantitative serology models that improve on these interval-censored uniform distributions have the potential to provide better estimates of the time of infection or seroconversion, which could dramatically improve estimates of epidemiological quantities such as incidence. As the potential error on this estimate can be large (up to several months), improved estimates of infection or seroconversion time obtained through quantitative serology would likely result in dramatic reductions in incidence estimation error.

Here, we present a general approach for modeling antibody dynamics when sampling is sparse and infection times are unknown (Figure 1). The approach uses hierarchical Bayesian MCMC inference to integrate different sources of information about model parameters, with full consideration and propagation of uncertainty, and accounting for population heterogeneity in serologic parameters. Additionally, we show how the simultaneous integration of the dynamics of additional biomarkers can lead to synergistic improvements in parameter fitting and infection time estimation. We apply this approach to Channel Island foxes (*Urocyon littoralis*) infected with *Leptospira interrogans* serovar Pomona. The framework presented here provides a way to estimate infection times by modeling biomarker dynamics even in the absence of experimental data, which we hope will stimulate more widespread use of quantitative serology in disease ecology.

**Figure 1.**
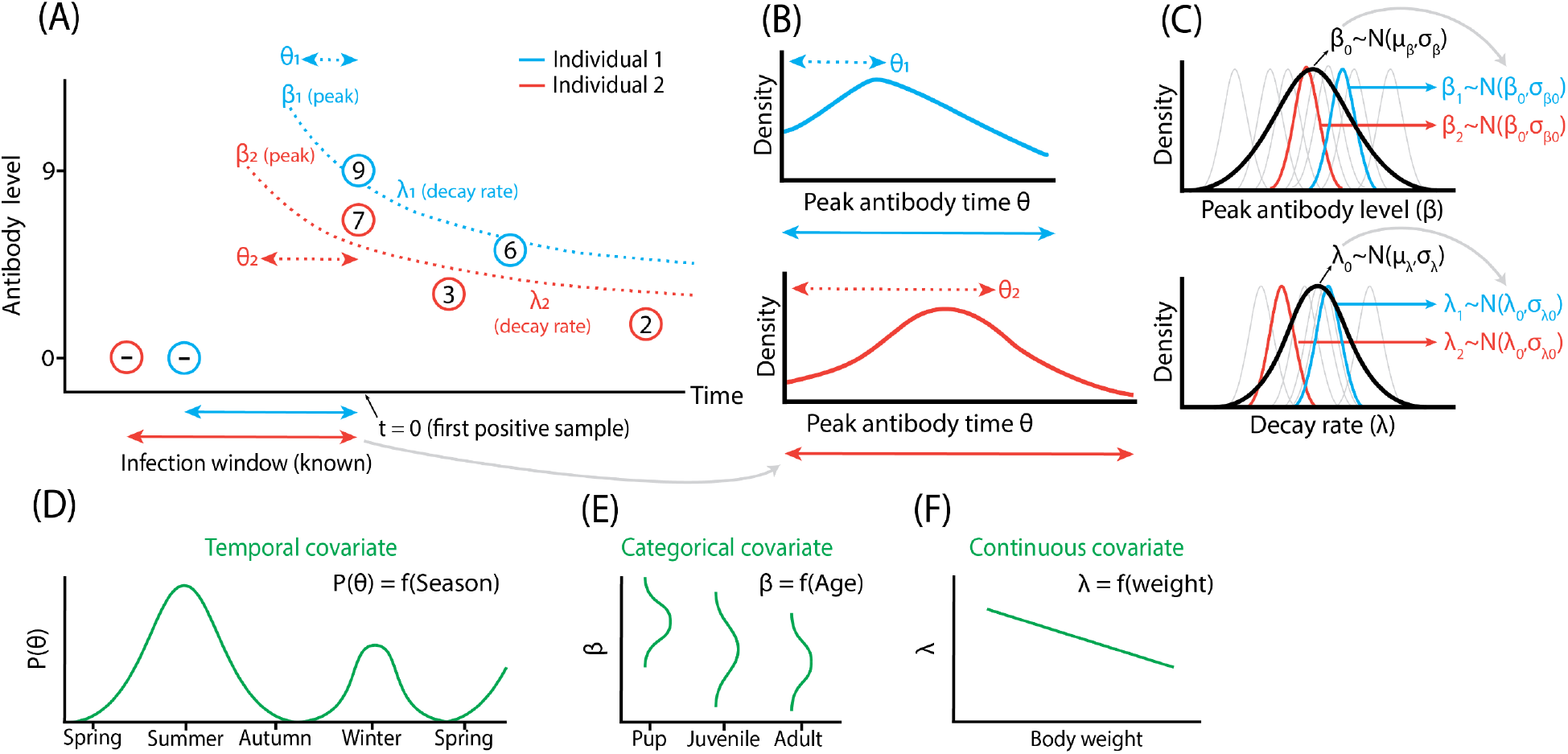
Hierarchical Bayesian modeling of biomarker data to estimate infection times. Bayesian inference offers a framework to use multiple sources of information to construct biomarker models and estimate individual time of infection. Panel (A) illustrates observed log_2_ antibody level data (circled numbers) for two individuals that are used to estimate model parameters *β* (peak antibody level) and *λ* (decay rate), with the ultimate goal of estimating time of infection for each individual *i* (which is determined by the peak antibody time *θ_i_*, measured relative to the time of the first positive sample) Dotted lines show possible unobserved antibody trajectories. Intervals between the last negative and first positive samples can be used as prior information to bound possible peak antibody times *θ_i_* (Panel B: posterior probabilities indicating the most likely peak antibody times). Model parameters can be estimated at the individual level (*β_i_* and *λ_i_*), while simultaneously estimating the mean and variation of these parameters at the population level (*β*_0_, *σ*_*β*0_, *λ*_0_, *σ*_*λ*0_) in a hierarchical way (C). When available (not in our study), other types of information can be used to improve estimates of the different model parameters, e.g. seasonal fluctuations in infection risk provide information about *θ_i_* (D), while age-dependent infection risk (E) or a continuous covariate such as body weight (F) can provide information about *β_i_* or *λ_i_*.

## Methods

### Data collection and sampling

Serum samples from Channel Island foxes were collected on Santa Rosa Island (California, USA) as part of a breeding, reintroduction and monitoring program (Coonan et al., 2015). Sampling was done annually between July and February, from 2004 through 2019. Foxes were trapped using Tomahawk Live Traps (Tomahawk Live Trap, Tomahawk, WI, USA; 0.66m × 0.23 m × 0.23m). Upon capture each individual was scanned for a passive integrated transponder (PIT) tag to uniquely identify it. If the animal was a new capture, a PIT tag was injected. Foxes were weighed, assessed for overall condition, age class, body condition and parasite load. Samples were collected during capture, taking up to 10ml of blood using a 22-gauge 1 inch needle. Blood was kept cold until centrifuging 3-5 hours later, after which serum samples were frozen. Samples were tested for antibodies against *Leptospira* using the microscopic agglutination test (MAT). Samples prior to 2016 were tested at the Centers for Disease Control (CDC; Atlanta, Georgia, USA) and samples from 2016 and later were tested at the Animal Health Diagnostic Center (Ithaca, New York, USA). All samples at both labs were titrated to endpoint titer against serovar Pomona. All samples tested at the CDC prior to 2013 were also titrated to endpoint titer for serovar Autumnalis. Antibody levels were log-transformed so that each unit change corresponds with a two-fold dilution step 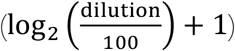. More details on trapping and sampling can be found in (Coonan, 2010). All trapping and sample collection was done by the National Park Service under a USFWS endangered and threatened species recovery permit for island foxes, *Urocyon littoralis*,TE-08267-2.

We used antibody levels against *Leptospira interrogans* serovar Pomona and serovar Autumnalis for antibody decay modelling and peak antibody time estimation. While the study population is known to be infected with serovar Pomona (Mummah, 2021), host antibodies can cross-react strongly, which means that MAT tests can also test positive for other serovars (Levett, 2003). As antibodies of foxes infected with serovar Pomona show a strong MAT signal for both serovar Pomona and serovar Autumnalis, we leveraged both data sources to improve model parameter fitting and infection time estimation.

### Candidate models

Prior to model fitting, candidate models of antibody decay had to be chosen based on preliminary exploration of the data. Generally, aspects to keep in mind when selecting candidate models are the possible shapes a function can have and the number of unknown function parameters. A model with more parameters results in higher flexibility but this can increase the risk of overfitting and reduce the model’s ability to predict new data (Bolker, 2008). Several functions have been used to model antibody decay, with the single (i.e. constant decay rate) and double (i.e. gradually decreasing decay rate) exponential functions being the most common (Boni et al., 2019; Teunis et al., 2016). When initial decay is significantly faster than later decay, alternative functions such as a power function can be used (Teunis et al., 2016). As shown in (Teunis et al., 2016), the power function may be particularly useful as it can accommodate a wide range of decay shapes with only two decay parameters. Additionally, the specifics of a power function support hypothesized underlying biological processes such as variation in the rate at which different sites in the body produce antibodies (Teunis et al., 2016). When empirical antibody kinetics do not resemble any existing functions, a flexible function such as a smoothed spline can be used (Borremans et al., 2016).

Based on initial data exploration and visualization we selected three candidate functions. Single exponential: *μ_i,t_* = *β_i_ e*^−*λ_i_*(*t*+*θ_i_*)^; double exponential: log (*μ_i,t_*) = log (*β_i_*) *e*^−*λ_i_*(*t*+*θ_i_*)^; power: 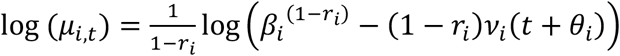; where *μ_i,t_* is the observed antibody level of individual *i* sampled at time *t*. Here, *t* is defined as the time since an individual’s first positive sample and *λ_i_* is the time of peak antibody level relative to the first observed positive sample of individual *i* (i.e. the number of days between an individual’s estimated time of peak antibody level and its first observed positive sample). *β_i_* and *λ_i_* are the peak antibody level and antibody decay rate of individual *i*. *r_i_* and *v_i_* are the shape and scale parameters of the power function. Functions were fitted to both Pomona and Autumnalis serovars, not considering mixed functions (e.g., a single exponential for Pomona and a double exponential for Autumnalis) based on the assumption that serovars will exhibit the same underlying decay process.

Note that we estimated the peak antibody level time and did not attempt to model the preceding period during which antibody levels increase, which is a limitation imposed by the low temporal resolution of our data relative to the duration of the increase period, which is not well known but is likely somewhere between 1 and 4 weeks (Langston and Heuter, 2003; Levett, 2001). In situations where data do allow quantification of the increase period, the increase and decrease phases are typically modeled as two different functions connected at the peak antibody level time (de Graaf et al., 2014; Teunis et al., 2016).

### Bayesian MCMC model fitting

Model fitting was done using a Bayesian Markov Chain Monte Carlo (MCMC) approach, as implemented in the software rJAGS (Plummer, 2019). A log-normal error distribution was assumed for antibody levels. Six parallel chains were run for 60,000 iterations, assessing chain convergence visually and with the Gelman-Rubin convergence diagnostic (Brooks and Gelman, 1998). Following a burn-in period of 10,000 iterations, posterior estimates were calculated for the last 50,000 iterations.

A key advantage of using a Bayesian approach for modelling antibody decay with unknown times of infection is the explicit incorporation of prior information as informative prior distributions for parameters. Here, we used informative priors for time of infection (implemented via the peak antibody level time, *θ_i_*) and peak antibody level *β_i_*, as described below. We further aimed to capture the biological variation in the model parameters across the population, to derive individual estimates as well as estimates of the mean and variation at the population level. This was possible by extending the Bayesian framework to a hierarchical structure, where the population-level parameters (now called hyperparameters) were estimated explicitly, and the individual-level parameters were drawn from these population-level distributions (Gelman and Hill, 2007). This meant that individual-level parameters were modeled using prior distributions 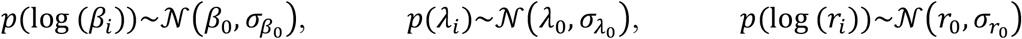 and 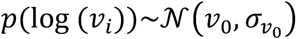, where *β*_0_, *σ*_*β*_0__, *λ*_0_, *σ*_*λ*_0__, *r*_0_ and *σ*_*r*_0__ are the hyperparameters of the hierarchical model: population-level means and standard deviations (sd) of peak antibody level (mean *β*_0_, sd *σ*_*β*_0__), exponential decay rate (mean *λ*_0_, sd *σ*_*v*_0__) and power function shape (mean *r*_0_, sd *σ*_*r*_0__) and scale (mean *v*_0_, sd *σ*_*v*_0__). Each of these hyperparameters had its own (hyper-)prior distribution (Table S1).

Parameter estimation and overall model performance can be greatly improved by combining data from multiple biomarkers or other covariates such as age, incidence seasonality and other infection risk factors (Borremans et al., 2016). As an example, we implemented an additional biomarker: antibody levels against *L. interrogans* serovar Autumnalis. This was possible using a joint-likelihood approach within the hierarchical Bayesian framework (Isaac et al., 2020). This is a simple extension of single-biomarker fitting, where two separate models (in this case, one for serovar Pomona and one for serovar Autumnalis) are fitted simultaneously, with distinct values allowed for all parameters except the peak antibody time *θ_i_*, which is constrained to be the same for both biomarkers. This approach increases the likelihood of accepting parameter values that are supported by the different biomarker datasets, and can result in more precise posterior estimates for *θ_i_*. Last, because Pomona and Autumnalis antibody levels are correlated, this potential correlation was implemented in the model by using a multivariate normal distribution for both peak antibody parameters.

Because samples were processed at two different labs over the 16 years of the field study, an additive lab effect parameter was added to the model. This allows for antibody levels of the labs to differ systematically.

The JAGS code used for model fitting has been provided as Supplementary Information (Section 8) and in the supplementary Rmarkdown code. Posterior 95% credible intervals (CrI) were calculated as highest density intervals using the function ‘dens’ of R package HDInterval (Meredith and Kruschke, 2018).

### Prior distribution of peak antibody time

Peak antibody time *θ_i_* was bounded by the interval between the most recent negative sample (a negative test result or birth date) and the first positive sample (the infection window). When available for an individual, this information was incorporated as a uniform prior distribution for *θ_i_* with minimum *θ_min_* and maximum 0: *p*(*θ_i_*)~*U*(*θ_min_*, 0). Alternative data to inform *θ_i_* include age (birth date), average lifespan when age data are not available, known seasonality in infection risk, onset of clinical signs of disease, and any other variable that provides information about the possible timing of infection. The probability distribution translating this information to a prior distribution can assume any shape and is not restricted to a uniform distribution as used here for the bounded infection window. We emphasize again that peak antibody time *θ_i_* is used as a proxy for time of infection, since the time resolution of the dataset makes it highly unlikely that sufficient samples were taken in the period between infection and peak antibody time.

### Prior distribution of peak antibody level

Another source of information that was used to improve model fitting is the distribution of peak antibody levels of recently infected foxes, which informs the population-level mean *β*_0_ and standard deviation *σ*_*β*_0__. This prior distribution was chosen by selecting a subset of foxes with relatively short infection windows, balancing the trade-off between sample size, which must be sufficiently large to provide a useful distribution, and recent infection time. We chose a maximum time of 250 days between the first positive and last negative sample as “recently infected”. Although this was still a large window, this was a limitation resulting from the frequency of sampling that is attainable in our field system and provides an opportunity to illustrate the strength of the approach under realistic field conditions. The limit of 250 days resulted in 54 out of 937 foxes that could be used to get an informed sense of the distribution of peak antibody levels at the population level. Normal distributions were fitted to the frequency distribution of the antibody levels of the first positive samples. Fitted means were increased slightly to account for antibody decay within the 250-day window, as were standard deviations to allow for additional uncertainty, resulting in prior distributions with mean 7 and sd 2.5 *log*_2_ dilutions for serovar Pomona and mean 7.5 and sd 3 *log*_2_ dilutions for serovar Autumnalis. Normal distributions were fitted using the fitdistr function in R package MASS (Venables and Ripley, 2002). More details are provided in Supplementary Information (Section 1).

### Model fitting

Model fitting was done using data from foxes that had at least 2 positive samples preceded by a negative one that determines the infection window. There were 34 out of 242 candidate foxes that exhibited signs of antibody boosting (possibly due to re-exposure to the pathogen, not necessarily leading to infection), defined here as an antibody level increase ≥ 2 log_2_ units between samples. Because the antibody model does not accommodate secondary increases in antibody level, these samples were removed from the dataset, starting from the sample preceding the secondary increase. These filtering rules were applied to both serovars, with samples exhibiting boosting only removed when the signal was present for serovar Pomona.

Model fits were compared using the leave-one-out cross-validation information criterion LOOIC (Vehtari et al., 2017) using R package loo (Vehtari et al., 2020), where lower values indicate a better fit, taking into account the number of parameters. Additionally, we used three measures that show the degree to which a model improves estimation of peak antibody time *θ_i_* relative to the uniformly distributed infection window bounded by the last negative and first positive sample. The first is “% reduction of the infection window” which is the percentage by which the size of the infection window was reduced when taking the 95% CrI of the posterior distribution as the new infection window. For example, if the original window size is 250 days (i.e. number of days between the most recent negative and first positive samples), and the model results in a posterior distribution of *θ_i_* for which the 95% CrI ranges from 200 to 20 days prior, the % reduction would be 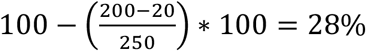. The second measure is the same as the first, but for 50% CrI ranges in order to better represent the estimates at the most concentrated area of the posterior. While these relative reduction measures are useful because they are easy to interpret, they do not consider that probabilities within the credible intervals are not equal, and some *θ_i_* will have a higher probability than others. To capture this, we used relative entropy (or Kullback-Leibler divergence) (Kullback and Leibler, 1951) as a second measure. Relative entropy (units = “bits”) quantifies the difference in information content between two distributions, which in this case are the uniform prior distribution (infection window) and the posterior distribution of *θ_i_*. Relative entropy 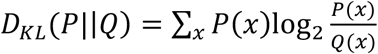, where *P*(*x*) and *Q*(*x*) are the posterior and prior distributions defined over the same range of values *x* (Burnham and Anderson, 2002). The values of *x* are individual-specific and adopt every possible value of *θ_i_* as determined by the uniform prior infection window. The higher the relative entropy value, the more information is present in the posterior distribution relative to the uniform prior.

### Sensitivity analysis using simulated data

To assess model performance given the limitations of our dataset, we simulated data mimicking the real dataset. Antibody levels were simulated for 75 individuals, where log peak antibody level and decay rate were randomly sampled from a normal distribution and 2 to 5 sampling events (random sample size) were simulated for sampling times up to 2,000 days after peak antibody level. Random noise was added to antibody levels to simulate real variation (see Supplementary Information Section 2 for details). For model fitting using simulated data, the peak antibody sample was excluded from the simulated dataset, again to mimic the real dataset. To test model sensitivity to different assumptions, parameter estimation was performed for multiple simulated datasets that were generated using a range of standard deviations for peak antibody level and decay rate. Details are provided in Supplementary Information (Section 2). We then tested how different combinations of peak antibody level variation and decay rate affect model performance, as faster decay and/or smaller variation in peak antibody level may constrain the possible peak antibody time window, which in turn would affect the precision of the peak antibody time *θ_i_* estimates.

### Software

All data preparation, analysis and plotting was done in R (R Core Team, 2019) using packages ggplot2 (Wickham, 2016), rjags (Plummer, 2019), ggridges (Wilke, 2020), dplyr (Wickham et al., 2019), patchwork (Pedersen, 2019), loo (Vehtari et al., 2020), R2OpenBUGS (Sturtz et al., 2005) and HDInterval (Meredith and Kruschke, 2018).

## Results

### Model fits to antibody decay data

Three different candidate models (single exponential, double exponential and power function) were fit to antibody data, and the best fits were observed for the two models (double exponential and power) that allow faster initial decay that slows with time since peak antibody level, for both serovar Pomona and Autumnalis. The double exponential model had the lowest LOOIC value (LOOIC values: single exponential = 5933, double exponential = 4705, power = 5757). All models were fit to the dataset including both serovar Pomona and Autumnalis, resulting in one overall LOOIC value but separate parameter estimatse for each serovar. The fitted functions using the population-level posterior means (reported below) are shown in Figure 2A. All further results will be shown for the double exponential model only. The fitted double exponential functions for serovars Pomona and Autumnalis are shown in Figure 2B with the observed data after adjusting time (x-axis) for each individual based on the estimated peak antibody time *θ_i_*. The mode (maximum posterior density) was used as the posterior estimate for *θ_i_* to accommodate the skewed posterior distributions on this parameter.

**Figure 2.**
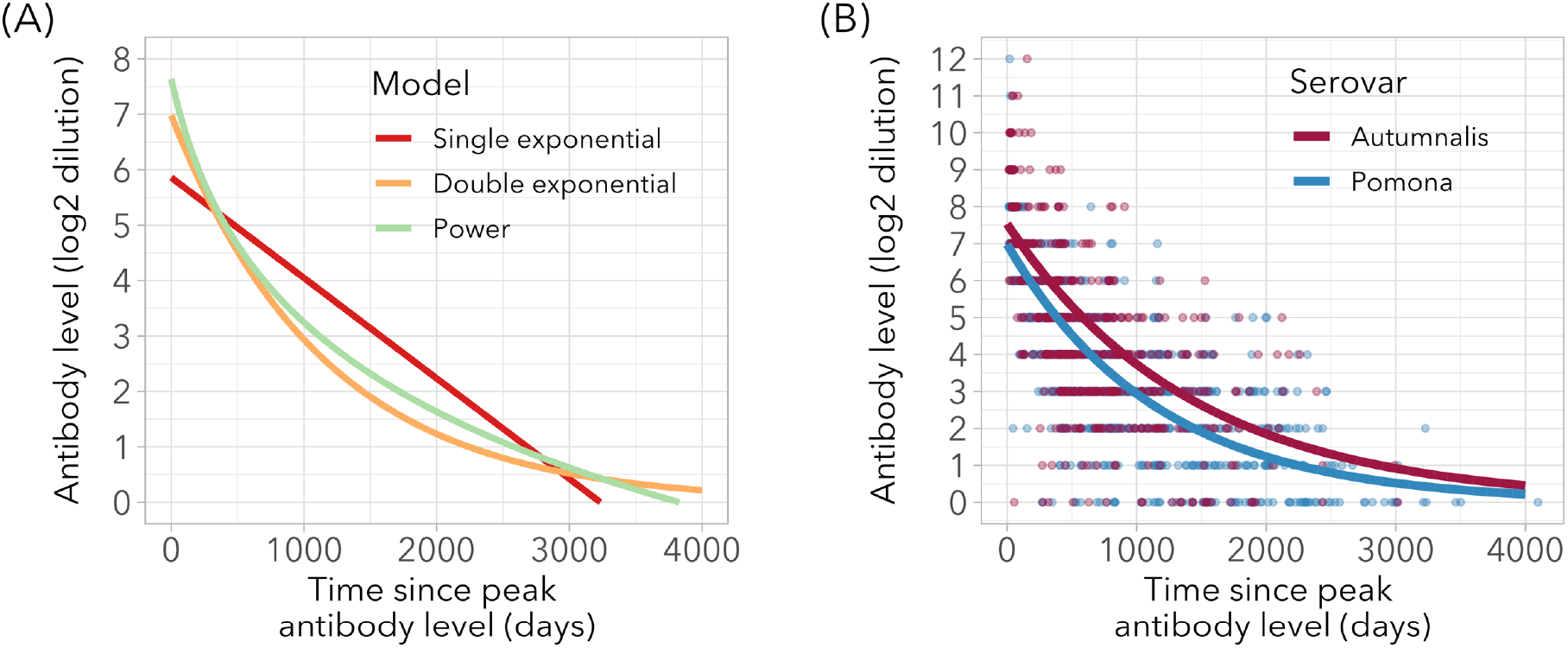
Candidate functions fitted to observed data. (A) Fitted functions for the three candidate models, using the posterior means for serovar Pomona peak antibody level and decay rate. (B) Fitted double exponential functions for serovars Pomona and Autumnalis, overlaid on observed data for 307 individuals after changing time since first positive sample to estimated time since peak antibody level.

### Posterior estimates of decay model parameters

This section provides the posterior estimates for all population-level parameters (results for each individual are shown in Figure 5). The population-level posterior means of the peak antibody level distribution parameters for serovar Pomona were 6.89 *log*_2_ units (95% CrI 6.61 to 7.15) for the mean and 2.00 *log*_2_ units (95% CrI 1.79 to 2.24) for the standard deviation. The estimated peak antibody level distribution for serovar Autumnalis was slightly higher, with a mean of 7.45 *log*_2_ units (95% CrI 7.11 to 7.88) and standard deviation of 2.08 *log*_2_ units (95% CrI 1.81 to 2.35) (Figure 3A).

**Figure 3.**
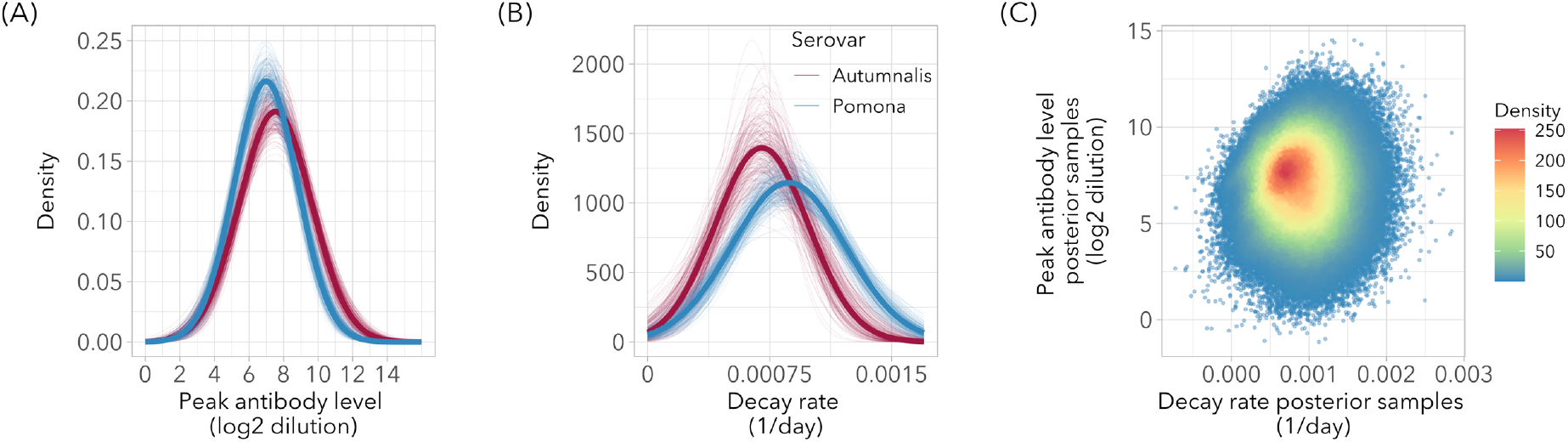
Estimated population-level distributions of peak antibody level (A) and decay rate (B), for serovars Pomona and Autumnalis. Bold lines are the distributions based on the posterior means of the mean and the standard deviation of each parameter. Thin lines are drawn from a random selection of 200 MCMC iterations, to show the magnitude of the variation around the posterior mean values. Panel (C) shows posterior samples for peak antibody level and decay rate, with colors depicting the density of points.

The population-level distribution of decay rate for serovar Pomona had a mean of 0.00086 *log*_2_ units/day (95% CrI 0.00079 to 0.00093) and standard deviation of 0.00035 *log*_2_ units/day (95% CrI 0.00028 to 0.00042). For serovar Autumnalis the mean was 0.00069 *log*_2_ units/day (95% CrI 0.00061 to 0.00078) and the standard deviation was 0.00028 *log*_2_ units/day (95% CrI 0.00021 to 0.00037) (Figure 3B). There was no statistically meaningful correlation between individual peak antibody level and decay rate (posterior mean of the linear regression slope: 90, 95% CrI −456 to 646; Figure 3C).

The mean percentage by which the individual prior infection windows were reduced after model fitting (using 50% credible intervals of the posterior distribution) was 61.3% (±0.6 SE, range 51.1 to 89.8%) when using the model that includes both serovars, and 61.2% (±0.4 SE, range 51.1 to 91.8%) for the model using serovar Pomona only (which contained more individuals, which explains the wider range). When using 95% credible intervals, mean window size reduction for the model including both serovars was 11.0% (±0.5 SE, range 5.0 to 53.4%) when using the model that includes both serovars, and 10.8% (±0.4 SE, range 5.0 to 66.7%) for the model using serovar Pomona only. Mean relative entropy values were 0.100 bits (±0.011 SE, range 0.006 to 1.185) and 0.107 bits (±0.014 SE, range 0.006 to 1.691).

### Correlates of model precision

We found strong individual variation in how much information the model was able to provide about peak antibody time *θ_i_*, with reductions in infection window size ranging from 51 to 92% (or 5% to 67% when using a 95% credible interval), and relative entropy values from 0.006 to 1.185 bits. To explore the factors underlying this variation in information gain, we tested the correlation between the reduction in an individual’s infection window (using the 95% CrI, which is highly correlated with the 50% CrI) and a number of variables: prior infection window size, number of samples, time range covered by the samples, estimated peak antibody level and
 estimated decay rate for serovar Pomona. Correlations were tested using linear models with a log-transformed infection window reduction outcome variable. We found that the model resulted in greater improvements in window size when the prior infection window was larger (effect size (rel. change) = 1.15, 95% CrI 1.10-1.20), when the estimated decay rate was higher (effect size (rel. change) = 1.21, 95% CrI 1.16-1.26), when the estimated peak antibody level was higher (effect size (rel. change) = 1.08, 95% CrI 1.03-1.13) and when the level of the first sample was higher (effect size (rel. change) = 1.06, 95% CrI 1.01-1.11). Model comparison using LOOIC showed the best fit for a regression model including a combination of decay rate, peak antibody level and infection window size, with decay rate occurring in all top models (Supplementary Information Section 3). In short, our method contributes more information when the early antibody levels are high (leading to high estimates of peak titer, and rapid decay) and when the possible infection window is long (meaning there is more room for improvement). This is illustrated in Figure 4, which shows the detailed output for two different individuals, while Figure 5 shows the posterior distributions of *θ_i_* for each individual.

**Figure 4.**
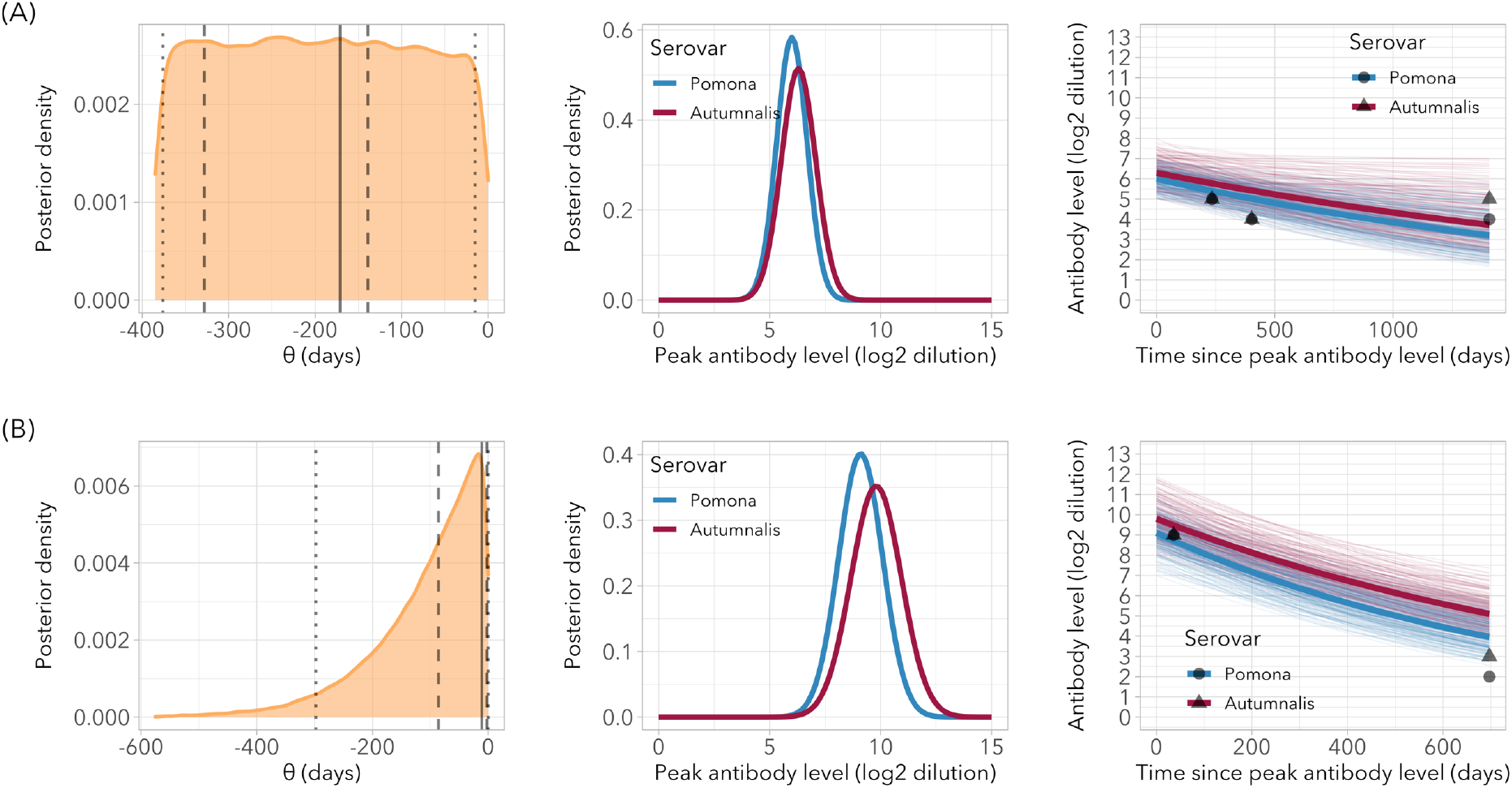
Posterior results for two individual foxes with low (A) and high (B) gain in information about peak antibody time *θ*. Left: posterior distribution of *θ*, the time between the first positive sample and the estimated peak antibody time, with maximum density (bold line) and credible intervals (95% = dotted line, 50% = dashed line). Center: posterior density for peak antibody level. Right: fitted functions using posterior estimates (bold lines) overlaid on 200 randomly selected iterations to show the distribution; black points are the observed samples.

**Figure 5.**
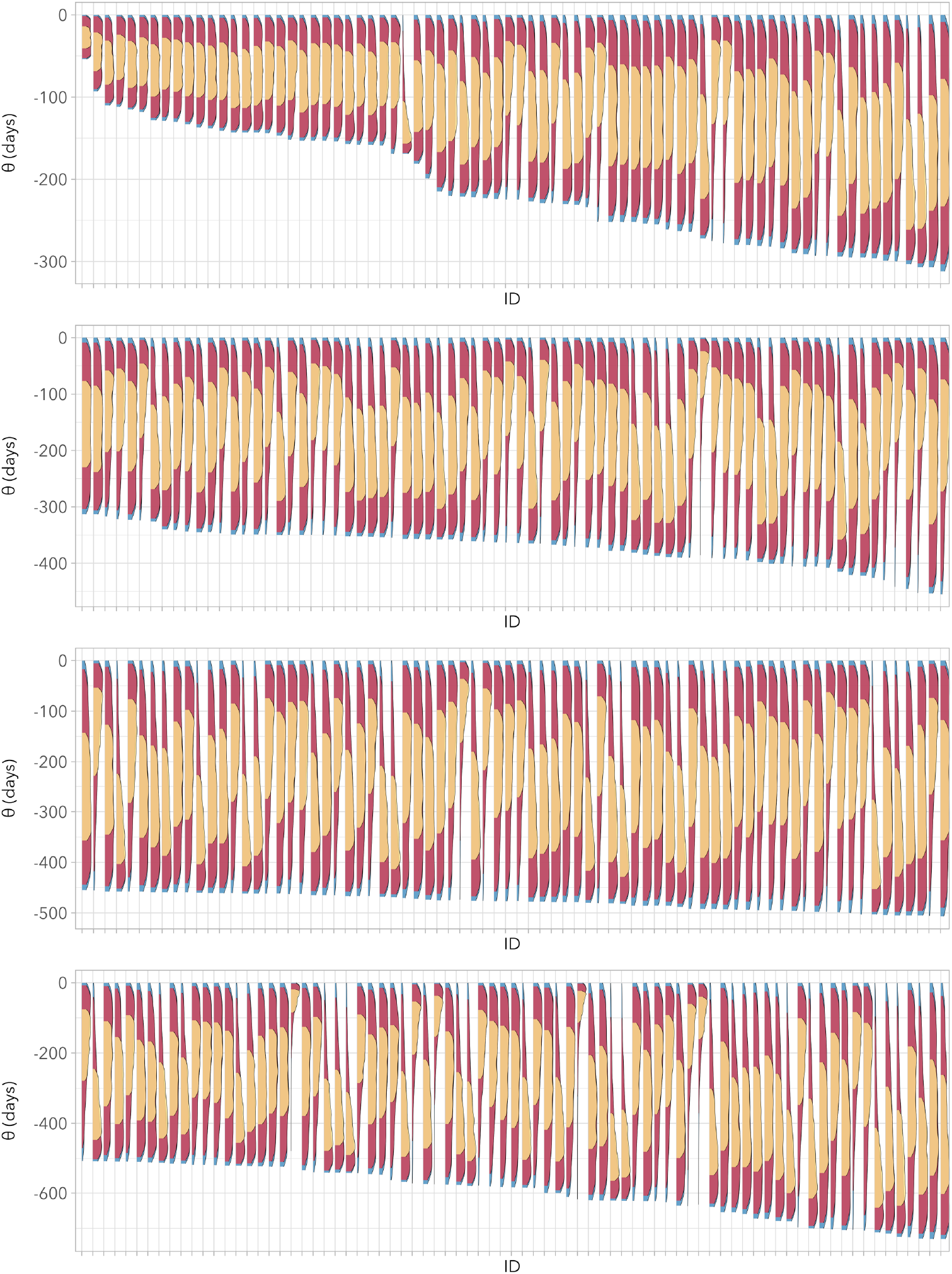
Posterior densities of peak antibody time *θ* for all foxes. Densities were plotted along the y-axis to enable showing all individuals, ordered along the x-axis by prior infection window size (full posterior density = blue, density within 95% CrI = red, density within 50% CrI = yellow).

### Model fitting performance using simulated data

The models fitted to simulated data (including added noise) were able to estimate the population-level parameters with accuracy and precision (*β*_0_, *σ*_*β*_0__, *λ*_0_ and *σ*_*λ*_0__; Figure S2) across a broad range of assumptions about peak antibody level mean and standard deviation, showing that the modeling approach works well. The estimation of parameter values at the individual level was good overall, but performance declined with more extreme values of the “true” individual peak antibody level (i.e. when the real simulated level was much lower or higher than the population mean; Figure S3 to S6).

Of broader relevance to biomarker dynamics in general, we found that model performance was strongly dependent on both decay rate and variation in peak antibody level (*σ*_*β*_0__). There was a strong positive correlation between decay rate and model performance (measured as infection window reduction), while there was a negative correlation between standard deviation of peak antibody level and model performance (Table 1). Figure S8 illustrates how the estimation of peak antibody time *θ_i_* will be more accurate with a faster decay and/or smaller variation in peak antibody values in the population.

**Table 1.** Effect of peak antibody level and decay rate on infection window estimation. Gained information on peak antibody time relative to the prior knowledge, a uniform interval bound by the last negative and first positive samples. Values show the percentage by which the infection window size was reduced, where the 95% CrI of the posterior distribution was taken as the new window. Results were similar when using relative entropy (Table S5)

| Peak<br>Ab SD<br>(log<sub>2</sub><br>units) | Infection window reduction (%) |  |  |  |
| --- | --- | --- | --- | --- |
|  | Decay rate (log<sub>2</sub> units/day) |  |  |  |
|  | 0.005 | 0.001 | 0.0005 | 0.0001 |
| 0.5 | 49 | 52 | 39 | 10 |
| 1.0 | 48 | 40 | 27 | 8 |
| 2.0 | 41 | 30 | 17 | 7 |
| 3.0 | 37 | 21 | 13 | 6 |

## Discussion

One of the outstanding challenges in quantitative serology is how to estimate time of infection from antibody levels in the absence of experimental data (Borremans et al., 2016; Pepin et al., 2017). By integrating data from different sources using a Bayesian approach, we were able to estimate time of infection and model antibody decay despite highly imprecise knowledge about when individuals were infected. Two key sources of prior information were used in the model. The first is information about possible peak antibody levels provided by a subset of foxes known to have been infected relatively recently. The second is the infection window, an interval that bounds each individual’s possible times at which antibody levels must have peaked, defined by a negative sample preceding the first positive one. Additionally, the Bayesian approach enabled leveraging data from multiple biomarkers, in this case antibody level data on *Leptospira interrogans* serovar Autumnalis in addition to serovar Pomona.

Model fit statistics showed a clear preference for the two models in which antibody decay decelerates with time, and of those the more parsimonious double exponential model received the highest level of support. Using the posterior means for the double exponential model, the decelerating decay when starting at a Pomona antibody dilution 1:6400 would translate to 198 days to decay to 1:3200, 431 days to 1:1600, 718 days to 1:800, and 2495 days to 1:100. Initially rapid antibody decay followed by a slow phase seems to be relatively common, and has for example been observed for leptospirosis in California sea lions (Prager et al., 2020), *Bordetella pertussis* in humans (Teunis et al., 2016) and arenavirus in multimammate mice (Borremans et al., 2015). This phenomenon may be a consequence of the heterogeneous dynamics of antibodies produced at different sites and different cell populations in the body, at different rates and driven by different mechanisms (Andraud et al., 2012; Teunis et al., 2016; Traggiai et al., 2003).

There was a strong correlation between antibodies against serovars Pomona and Autumnalis, which was particularly strong for peak antibody levels. This meant that the information gained by the addition of data on serovar Autumnalis was limited, as that information was for the most part already provided by serovar Pomona. We indeed found that the reduction in infection window size achieved using our model was not much better when integrating data on both serovars versus only using serovar Pomona. This is a useful general insight into what to expect when considering incorporating multiple biomarkers for quantitative serology, and suggests that a useful strategy would be to prioritize biomarkers that do not correlate strongly. For instance, combining antibody data with data on the presence of a pathogen (or its genetic material) is likely to be much more informative than data on an antibody exhibiting the same dynamics, as shown for arenavirus infection in the rodent *Mastomys natalensis* (Borremans et al., 2016), hantavirus in *Peromyscus sp*. mice (Abbott et al., 1999) and influenza A in swine (Strelioff et al., 2013). Alternatively, biomarkers that reflect the state of disease (e.g. renal health) can add useful information (Prager et al., 2020).

We found that population-level estimates of peak antibody level and decay rate could be reproduced well in simulations, and while most individual-level estimates were close to the simulated values, some were not. Closer inspection showed that this happens when an individual’s peak antibody level was much lower or higher than average. Because the model assumes that individual levels are a sample of the population-level distribution, values closer to the mean will have a higher likelihood. This “shrinks” the individual estimates towards the population means, which is generally not problematic unless the real individual peak antibody value is much lower or higher. This is by definition rare when peak antibody levels or decay rates are assumed to be distributed normally; future work could consider hyper-parameter distributions with heavier tails.

The degree to which the model was able to improve individual infection windows correlated strongly with a number of individual variables. The model yielded greater reductions in infection window with higher decay rate, higher peak antibody level, higher level of the first positive sample and greater size of the prior infection window size. The latter can be explained by the fact that when the prior infection window is longer, larger improvements are possible given the fact that decay rate and peak antibody level are restricted by the population-level parameter values. Higher decay rates on the other hand mean that the information conveyed by antibody levels will be more localized in time, thus resulting in better estimates of peak antibody time (Table 1 and Figure S8). The effect of a higher level of peak antibody or first positive sample results from the fact that decay is faster at higher levels, and that the closer the level of the first positive sample is to the high end of the peak antibody level distribution, the smaller the possible infection window. Last, as expected we observed in both the data and the simulations that a broader distribution of possible peak antibody levels directly translates into a broader distribution of possible peak antibody times, which means that lower individual variation will result in better estimates of time of infection (Table 1).

These characteristics related to model performance are intrinsic to a pathogen-host system, and will determine the upper level of performance of any model. For our system, the model yielded reductions in infection window size ranging from modest (51% using 50% credible intervals, or 5% using 95% intervals) to impressive (92% or 67%, respectively); these reductions were obtained despite the relatively slow decay of antibodies against *L. interrogans* serovar Pomona in Channel Island foxes as well as considerable variation in peak antibody level. Indeed, the estimated decay rate of 0.0009 *log*_2_ units/day results in a window of 320 days between antibody levels 8 and 6 *log*_2_ units (assuming a peak level of 11). For comparison, with decay rates of 0.005 and 0.01 *log*_2_units/day would result in windows of 58 and 29 days, respectively. This means that for other systems such as *L. interrogans* serovar Pomona infection in California sea lions (*Zalophus californianus*) where antibody decay can be as fast as 0.058 (Prager et al., 2020), a similar model would be expected to yield considerably more precise estimates of peak antibody time.

Study systems where sampling is seasonal, as is the case for many wildlife systems, can benefit particularly from this method. Island fox field work and sampling is highly seasonal, resulting in gaps of several months without any data that create a major challenge for interpreting and modeling temporal disease dynamics. Our method however provides estimates of time of infection for any time, including periods without sampling, thus providing a much better insight into disease dynamics over time. Additionally, once the model has been trained using “high-quality” data, it can be applied to individuals with any number of positive samples, even a single one as is the case for cross-sectional sampling (Figures S11-13). There is even no requirement for a preceding negative sample or other information such as age to delineate an infection window, as in such cases an arbitrary window can be used that is longer than any possible window (e.g. the maximum possible lifespan).

Regardless of system characteristics and data quality, there will always be a certain amount of uncertainty due to biological variation and observational noise. Another major benefit of using a Bayesian MCMC approach is the fact that it explicitly integrates all sources of variation and uncertainty, resulting in a probability distribution of all parameter estimates instead of a single point estimate with a standard error when using frequentist methods. All known sources of uncertainty are therefore acknowledged in the posterior distributions. This, for example, provides a useful platform for principled averaging over possible times of infection, as shown in related work in which individual times of infection were used for survival models of *Leptospira* infection in Channel Island foxes (Mummah, 2021).

Another advantage of the Bayesian approach is that it enables the use of information from widely varying sources (Figure 1). As an example, we have shown here that it is possible to incorporate an additional biomarker (antibodies against *L. interrogans* serovar Autumnalis). Other candidate data sources include individual age, sex or body condition. For example, we tested whether any of these three variables correlated with the antibody level of the first positive sample of recently infected individuals (Supplementary Information Section 5). In our case none of the correlations were statistically meaningful (all 95% credible intervals contained 0), but if such a correlation existed then it could be included in the model as prior information for peak antibody level. The strength of the combined use of different sources of information can be seen for the estimation of the time of infection in multimammate mice infected with an arenavirus, where incorporating antibody level, virus RNA presence, individual age and encounter probability resulted in large improvements in model performance (Borremans et al., 2016). It is important to note that while a Bayesian approach has many advantages, as with all models care must be taken on how they are constructed and interpreted. For example, combining data sources must be done carefully to avoid spurious results, while the use of informative priors can influence the analysis and must also be done with care.

Once a model to estimate infection time (or peak antibody time) has been constructed and parameterized using an informative dataset of individuals, it can be applied to further data sets of lower quality or temporal resolution. These data need not include a negative sample preceding the first positive one, and can even consist of a single result per individual (see Supplementary Information Section 7). This enables the application of these quantitative serology methods to cross-sectional data, providing a powerful tool for wildlife disease ecologists. A further advantage of the Bayesian approach is that it is trivial to improve the model when new informative data become available, as the same model can be fitted using the expanded dataset or in a sequential manner using the posteriors as priors (Bolker, 2008).

Quantitative serology is a growing field with major potential (Gilbert et al., 2013; Held et al., 2019; Pepin et al., 2017; Teunis et al., 2012; Wilber et al., 2020), but its broad adoption has been relatively limited. While this may partly be due to a lack of exposure, it is likely that a number of outstanding challenges are keeping scientists from applying these methods. One of these challenges, especially for wildlife systems with sparse sampling, is the difficulty in constructing biomarker models when there are no experimental data mapping out longitudinal patterns, and when infection times are unknown. As most biomarker models are specific to a host-pathogen system, they need to be constructed and parameterized for each system before they can be used to estimate time of infection. Here, we have provided a complete approach to constructing biomarker models entirely from field data where sampling points are sparse and infection times unknown, using a hierarchical Bayesian framework that enables leveraging multiple sources of information and multiple types of data (Figure 1). This addresses a major challenge in quantitative serology and time-of-infection estimation that has been one of the key barriers for application to wildlife systems. This work opens new frontiers for quantitative analysis of infectious disease dynamics, including incidence reconstruction with proper error propagation, anamnestic immune response following re-exposure (boosting), dealing with antibody cross-reactivity, integration of infection time estimation and transmission modeling, and software development for easy integration of data sources and incidence reconstruction.

## Acknowledgments

BB was supported by the European Commission Horizon 2020 Marie Sklodowska-Curie Actions (grant no. 707840). JOL-S was supported by the Defense Advanced Research Projects Agency DARPA PREEMPT # D18AC00031 and the UCLA AIDS Institute and Charity Treks. JOL-S and KCP were supported by the U.S. National Science Foundation (DEB-1557022 and OCE-1335657), the Strategic Environmental Research and Development Program (SERDP, RC-2635) of the U.S. Department of Defense, and the Cooperative Ecosystem Studies Unit Cooperative Agreement #W9132T1920006. The authors would like to express their gratitude and respect for the tremendous work performed by all personnel involved in the field work. The findings and conclusions in this report are those of the author(s) and do not necessarily represent the official position of the Centers for Disease Control and Prevention. All code and data are available online at github.com/bennyborremans/antibody_decay_field_data.

