## Supplementary Information for "Inferring time of infection from field data using dynamic models of antibody decay"

# to

**Preprint**

This manuscript is a preprint and has not been peer-reviewed. It is currently under review at a peer-reviewed scientific journal. We welcome any feedback, through the comments section on the bioRxiv website or by emailing the corresponding author.

All code and data are available online at github.com/bennyborremans/antibody_decay_field_data.

### 1. Prior distribution of peak antibody level, and hyperpriors

A subset of recently negative individuals was used to provide information about what the possible distribution of peak antibody levels could be. Antibody levels were assumed to have peaked during the interval between the last negative timepoint preceding the first positive one. Here, a negative timepoint could be either an earlier negative sample or the animal's birth date. After data exploration, a subset of individuals was selected for which the maximum time between the last negative and first positive timepoints was 250 days. This resulted in a dataset of 54 individuals. Although a period of 250 days is still a relatively long interval, selecting a smaller interval would have resulted in a much smaller dataset due to the seasonality of sampling in our system. To account for this extended interval we used a prior distribution with a slightly larger mean than the distribution fitted to the observed data, but with a broader standard deviation. This resulted in a prior distribution that is weakly informative, thus allowing a large influence of the data on the posterior likelihood. Figure S1 shows the distribution of observed antibody levels and the fitted normal distributions. The distribution was fitted using the function fitdistr of R package MASS (Venables and Ripley, 2002). Using simulated data with characteristics similar to those of the real data, we found that the effect of the prior distribution specifications was not large, and that the population mean peak antibody level could be estimated well (see below).

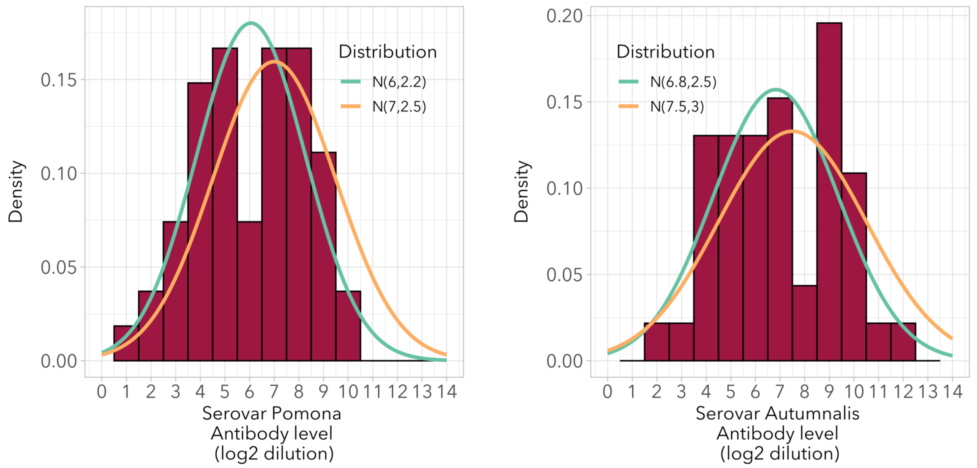


Figure S1. Antibody levels for a subset of individual foxes that tested negative within at most 250 days (left: serovar Pomona, right: serovar Autumnalis). Fitted distributions are shown in teal, and distributions used as priors are shown in orange.

### Hyperpriors

Table S1. Hyperpriors for the different parameters.

| Parameter | Prior distribution |
|---|---|
| $\boldsymbol{\beta}_{0,Pomona}$, $\boldsymbol{\beta}_{0,Autumnalis}$ | $p(\log(\boldsymbol{\beta_0})) \sim \mathcal{N}(\boldsymbol{\beta_0^h}, \boldsymbol{\Sigma_{\beta 0}^h})$,<br>multivariate normal distribution with mean vector $\boldsymbol{\beta_0^h} = \begin{bmatrix} 7 \\ 7.5 \end{bmatrix}$ and covariance matrix $\boldsymbol{\sigma_{\beta 0}^h} = \begin{bmatrix} 2 & 0 \\ 0 & 2 \end{bmatrix}$. |
| $\boldsymbol{\sigma}_{\beta_{0,Pomona}}$, $\boldsymbol{\sigma}_{\beta_{0,Autumnalis}}$ | $p(\boldsymbol{\sigma_{\beta 0}}) \sim Wishart(\boldsymbol{\mathcal{V}}, df)$,<br>Wishart distribution with scale matrix $\boldsymbol{\mathcal{V}} = \begin{bmatrix} 2.5 & 0 \\ 0 & 3 \end{bmatrix}$ and degrees of freedom $df = 2$. |
| $\boldsymbol{\lambda_0}$ | $p(\lambda_0) \sim Gamma(1{,}20)$,<br>Gamma distribution with the same parameters for serovars Pomona and Autumnalis. |
| $\boldsymbol{\sigma_{\lambda_0}}$ | $p(\sigma_{\lambda 0}) \sim Gamma(2{,}3)$,<br>Gamma distribution with the same parameters for serovars Pomona and Autumnalis. |
| $\boldsymbol{r_0}$ | $p(r_0) \sim \mathcal{N}(\log(0.5), 0.1)$,<br>Normal distribution with the same parameters for serovars Pomona and Autumnalis. |
| $\boldsymbol{v_0}$ | $p(v_0) \sim \mathcal{N}(\log(0.0005), 1)$,<br>Normal distribution with the same parameters for serovars Pomona and Autumnalis. |

### 2. Simulations and sensitivity analysis

The model fitting approach was tested using simulated data (with characteristics similar to those of the real data) for 75 individuals, using the double exponential function (see main text). For each individual a peak antibody level representing serovar Pomona was randomly generated from a certain normal distribution that depended on the simulated scenario (see below). A second peak antibody level representing serovar Autumnalis was generated from a different distribution of which the mean was 0.84 $log_2$ (with a standard deviation of 0.5 to add realistic variation) units higher. Next, a decay rate was randomly generated from a certain distribution (see below), where the mean of the distribution for serovar Autumnalis was 20% higher than that for serovar Pomona. Using those parameters, between two and five samples (the number was randomly chosen with equal probability) were generated for each individual at random times after the peak antibody time, with a maximum time of 2000 days. A negative sample preceding the first positive sample was also generated to inform the maximum infection window size, where the time was generated from a uniform random distribution with a maximum of 500 days prior to the peak antibody time. Last, noise was randomly added to the data where each sample had a probability of 5% to decrease by one $log_2$ unit, 90% to remain the same, and 5% to increase by one $log_2$ unit. This represents plausible variation resulting from the microscopic agglutination assay (see main text) in two-fold dilution units. Each individual's peak time sample was removed from the dataset prior to model fitting, to represent reality.

Simulations were done for two scenarios:

1. **Effect of peak antibody level prior specification**. In order to test the effect of the specifications of the prior distribution of peak antibody level on the posterior estimate, we fitted the same simulated data using different prior specifications of peak antibody. Peak antibody levels (serovar Pomona) were generated from a normal distribution $\mathcal{N}(7,1)$, and decay rates from $\mathcal{N}(0.0008, 0.0002)$. Model parameters were fit for six prior distributions of peak antibody level, to assess the effect of the mean and the standard deviation on the posterior estimates: $\mathcal{N}(6,1)$, $\mathcal{N}(8,1)$, $\mathcal{N}(10,1)$, $\mathcal{N}(8,2)$, $\mathcal{N}(8,3)$, $\mathcal{N}(8,4)$. For all prior distributions the posterior estimates accurately estimated mean peak antibody level at the population level (Figure S2). Individual-level estimates of peak antibody time were good overall

(Figure S3 and S4), but when the true (i.e. simulated) peak antibody level was at the extremes of the population-level distribution the estimates became less accurate.

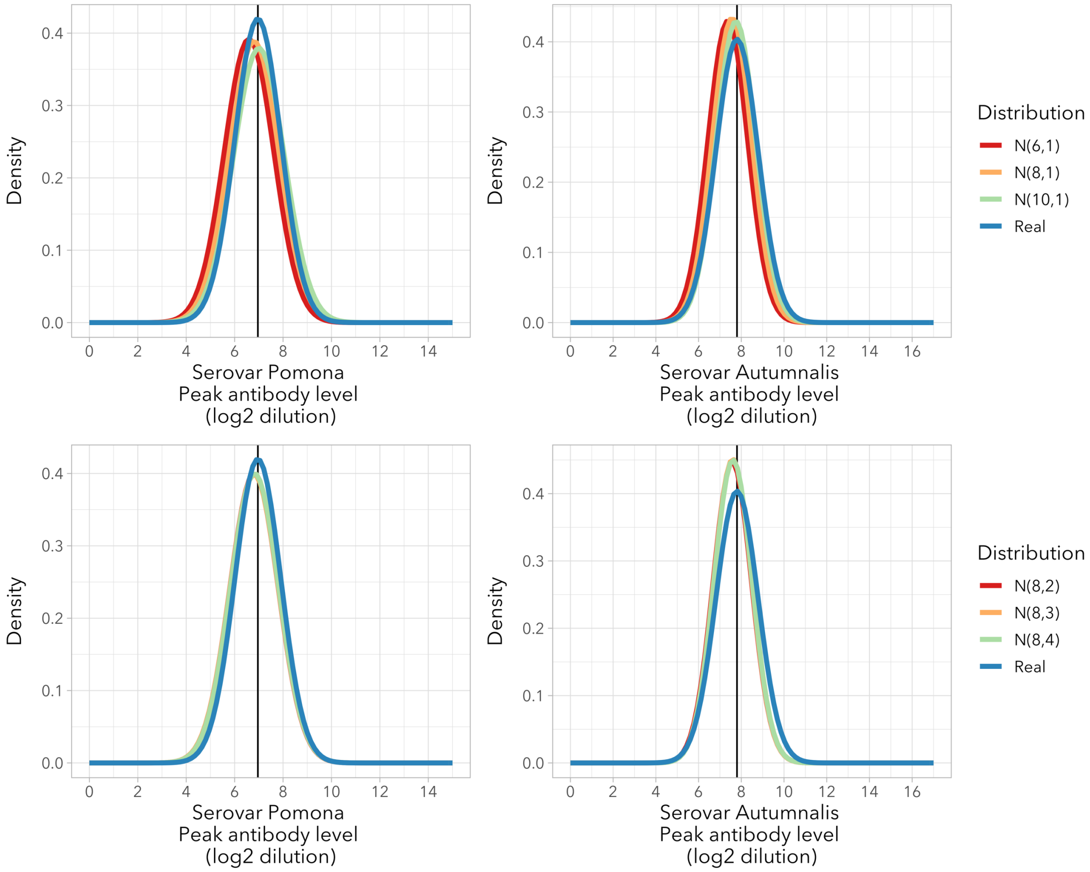


Figure S2. **Posterior estimates of peak antibody level for different prior distributions**. True = generated distributions with means 7 (SD 1) and 7.8 (SD 1) for serovars Pomona and Autumnalis, respectively.

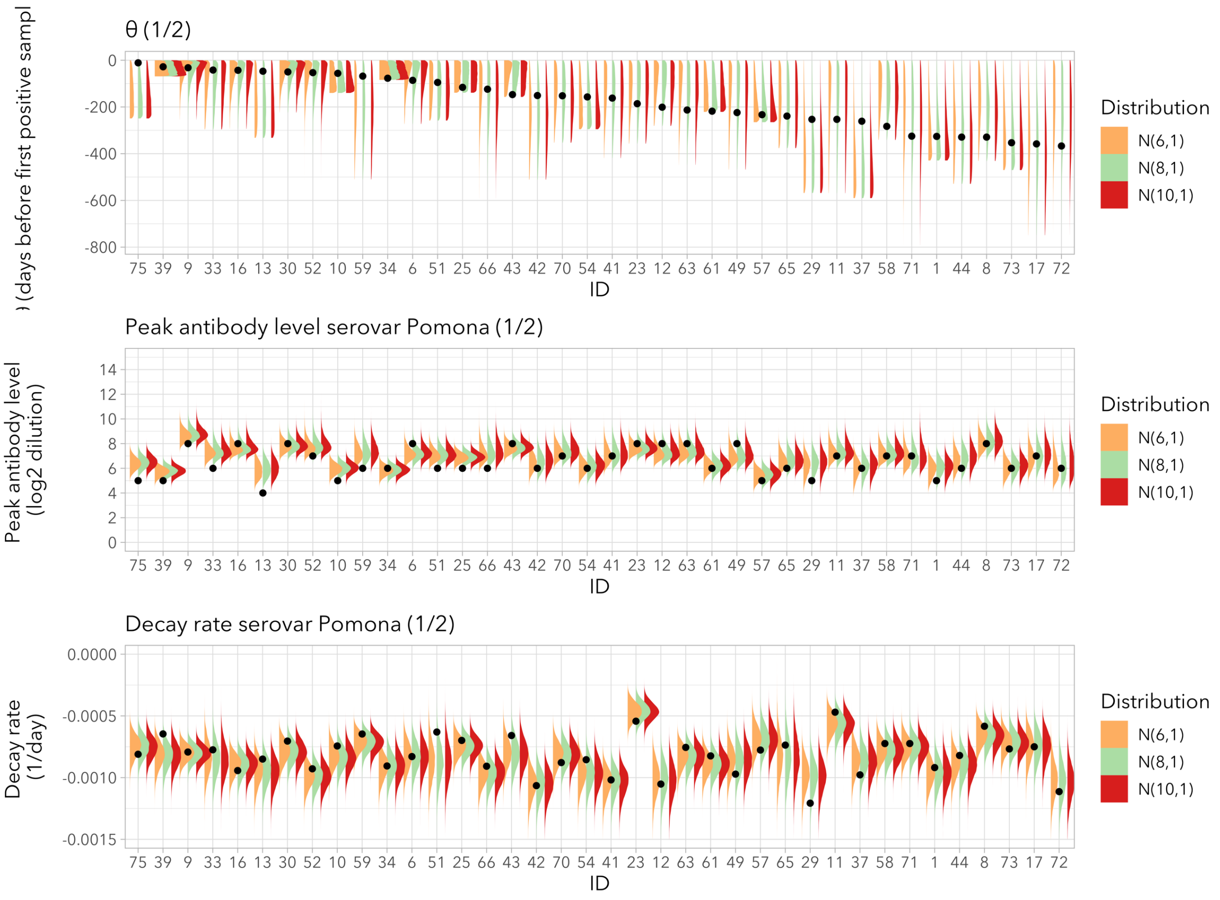


Figure S3. **Effect of prior peak antibody level means**. Posterior densities of peak antibody time, peak antibody level and decay rate for the first half of the 75 simulated individuals. Dots indicate the "real" simulated value, distributions are posterior densities flipped vertically.

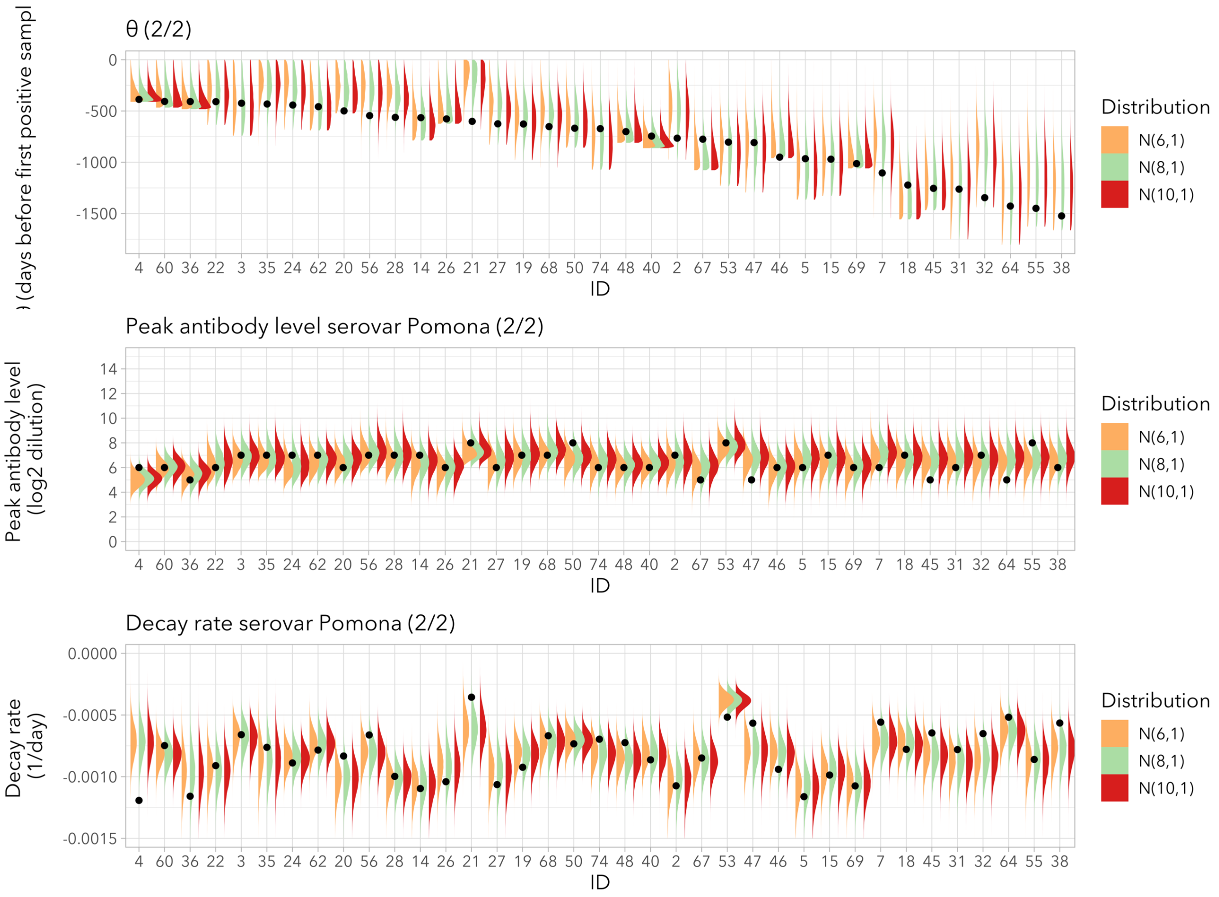


Figure S4. **Effect of prior peak antibody level means**. Posterior densities of peak antibody time, peak antibody level and decay rate for the second half of the 75 simulated individuals. Dots indicate the "real" simulated value, distributions are posterior densities flipped vertically.

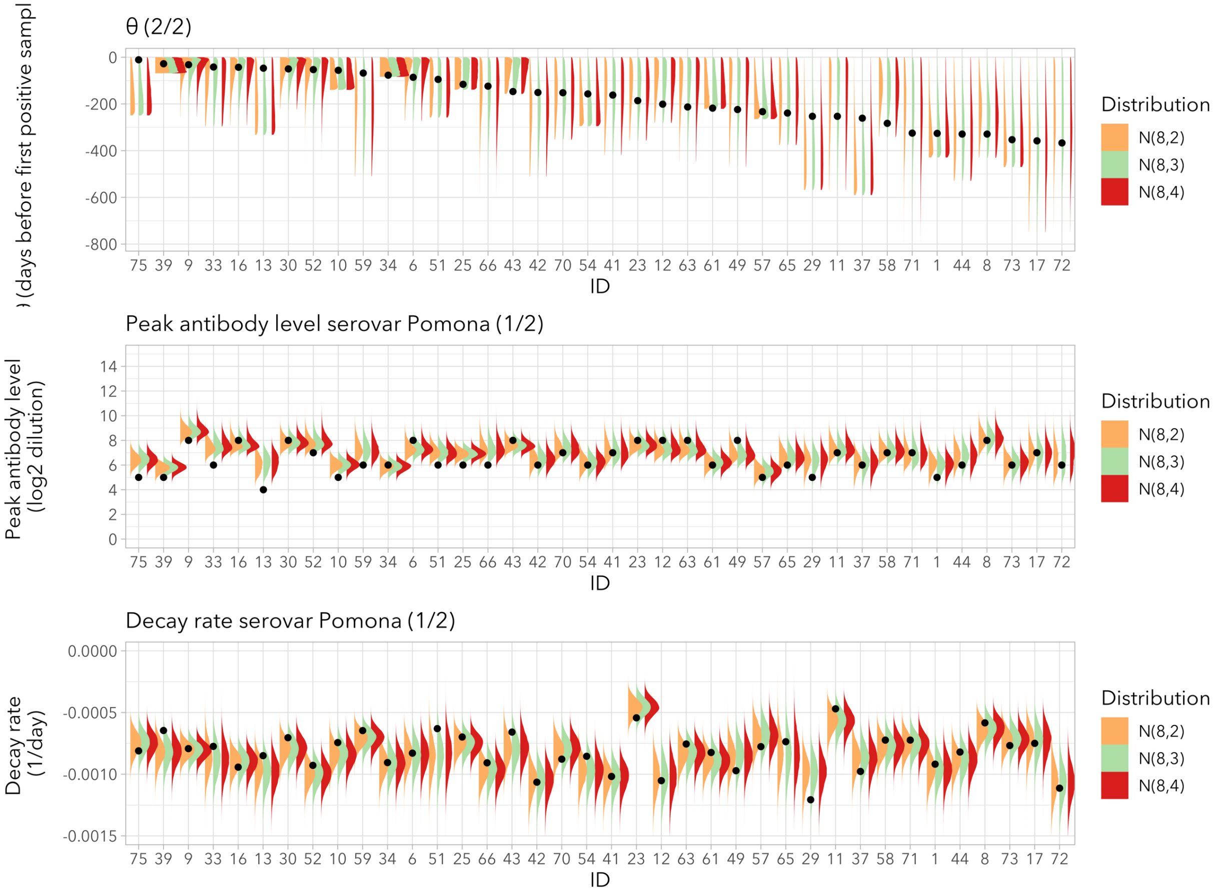


Figure S5. **Effect of prior peak antibody level standard deviations**. Posterior densities of peak antibody time, peak antibody level and decay rate for the first half of the 75 simulated individuals. Dots indicate the "real" simulated value, distributions are posterior densities flipped vertically.

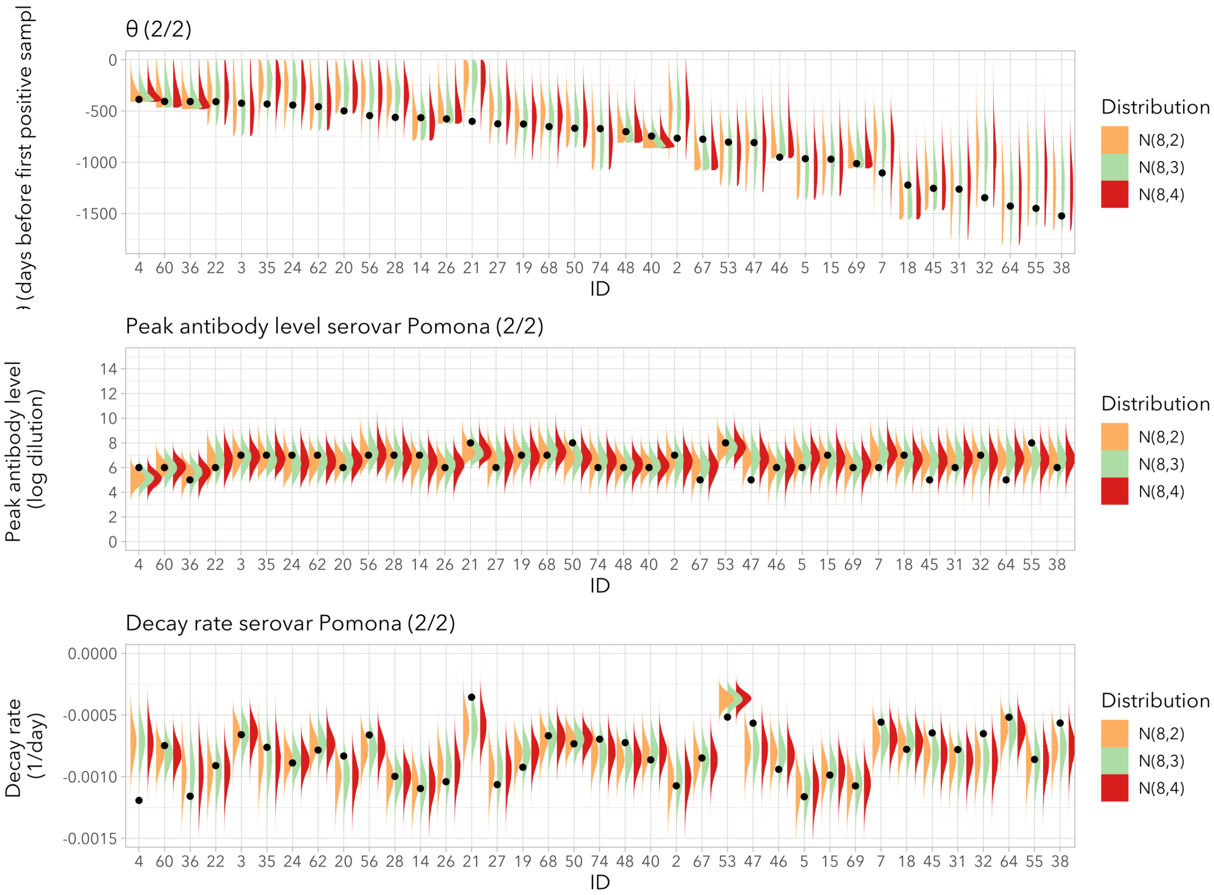


Figure S6. **Effect of prior peak antibody level standard deviations**. Posterior densities of peak antibody time, peak antibody level and decay rate for the second half of the 75 simulated individuals. Dots indicate the "real" simulated value, distributions are posterior densities flipped vertically.

2. **Effect of the magnitude of peak antibody level and decay rate**. Mean peak antibody level as well as decay rate are expected to have a strong effect on how well the models will be able to estimate peak antibody time. In order to quantify this, we fitted models to a series of datasets simulated using a range of peak antibody levels and decay rates. The effect of the different parameter values was then quantified as the mean reduction in the infection window. Peak antibody levels (serovar Pomona) were generated from distributions with mean 7 and standard deviations 0.5, 1, 2 and 3. Decay rates (serovar Pomona) were generated from normal distributions with means 0.005, 0.001, 0.0005, 0.0001 and standard

deviation 0.0002. Figure S7 shows antibody decay for the different decay rates. Model parameters were fit for each combination of peak antibody level and decay rate distributions. Model performance was clearly dependent on the characteristics of antibody decay, where smaller variation in peak antibody level and/or faster decay resulted in more accurate estimates of peak antibody time, and larger gains of the posterior distribution of peak antibody times relative to the prior uniform interval (Table S2).

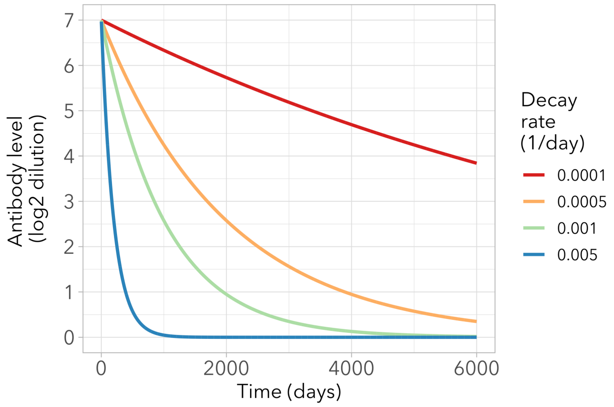


Figure S7. **Antibody decay curves for different decay rates**.

Last, in order to quantify the effect of decay rate on how well the method can be expected to estimate peak antibody time, we calculated the time needed to decay from antibody level 8 to 6 $\log_2$ dilution units for a range of decay rates (Figure S8), choosing these two levels because they are close to peak levels but at the mean to lower end which means they are quite common and relevant. A larger time window will result in a less precise estimate of peak antibody time, and Figure S8 clearly illustrates that decay rate will be a major determinant of how well a time-of-infection estimation method can be expected to perform.

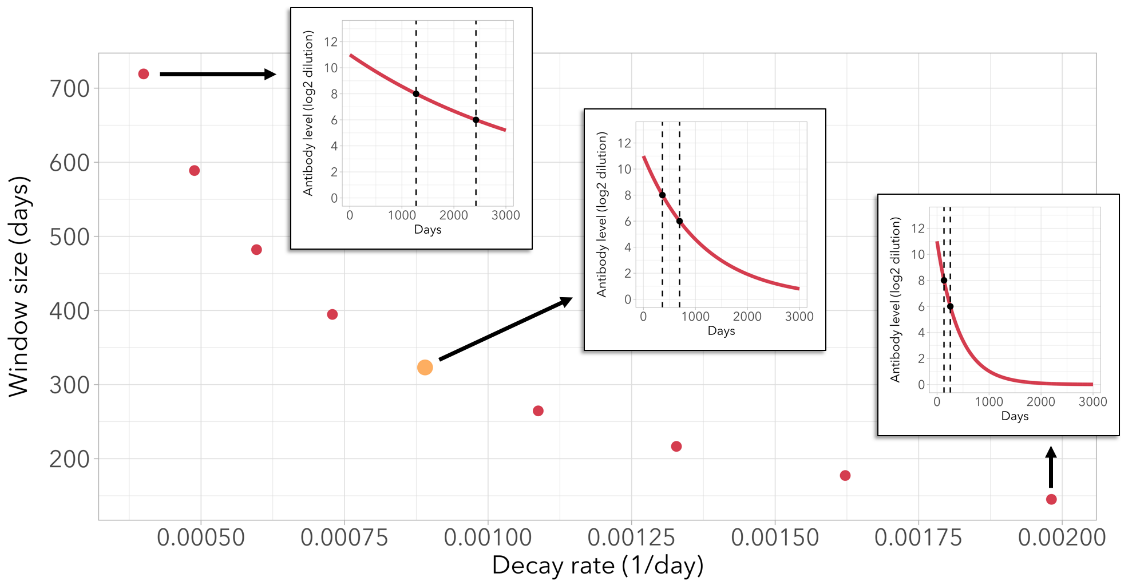


Figure S8. **The number of days (window size) it takes for antibodies to decay from 8 to 6 $\log_2$ dilution units, for a range of biologically realistic decay rates**. The yellow dot marks a decay rate close to the one estimated for serovar Pomona. Inset figures show the decay function for selected decay rates, with the window between antibody levels 8 and 6 indicated with dotted lines.

### 3. Predictors of model performance

We tested whether any variables correlated with a reduction in an individual's infection window size. Candidate variables were prior infection window size, number of samples, time range covered by the samples, estimated peak antibody level and estimated decay rate for serovar Pomona. Correlations were tested using linear models with a log-transformed interval reduction outcome variable. Results are shown in Table S2, Table S3 and Figure S9.

Table S2. Statistics for the correlations between outcome variable '% infection window reduction' and candidate variables, using a 95% credible interval for the infection window posterior. Effect sizes are exponentiated to get relative change and are shown with 95% credible intervals. LOOIC values are shown with standard errors.

| Variable | Effect size (exp) | Effect size l-95% | Effect size u-95% | LOOIC | LOOIC SE |
|---|---|---|---|---|---|
| **Prior infection window size** | 1.15 | 1.10 | 1.20 | 300 | 42 |
| **Number of samples** | 0.96 | 0.92 | 1.01 | 333 | 43 |
| **Time range of samples** | 0.99 | 0.95 | 1.04 | 335 | 43 |
| **Peak antibody level posterior mean** | 1.08 | 1.03 | 1.13 | 325 | 39 |
| **Decay rate posterior mean** | 1.21 | 1.16 | 1.26 | 262 | 42 |
| **Antibody level of the first positive sample** | 1.06 | 1.01 | 1.11 | 331 | 39 |

Table S3. LOOIC values for linear regression models including single and multiple variables fitted to outcome variable '% infection window reduction'. Sorted by LOOIC value, which are shown with standard errors.

| Model | LOOIC | LOOIC SE |
|---|---|---|
| **Peak level + decay rate + window size** | 159 | 32 |
| **All variables** | 161 | 32 |
| **Decay rate + window size** | 212 | 41 |
| **Decay rate + peak level** | 232 | 34 |
| **Peak level + decay rate + number of samples** | 234 | 34 |
| **Decay rate + first positive level** | 251 | 36 |
| **Decay rate posterior mean** | 262 | 42 |
| **Peak level + window size** | 284 | 36 |
| **Prior window size** | 300 | 42 |
| **Peak level posterior mean** | 325 | 39 |
| **Peak level + number of samples** | 326 | 39 |
| **Antibody level of the first positive sample** | 331 | 39 |
| **Number of samples** | 333 | 43 |
| **Time range of samples** | 335 | 43 |

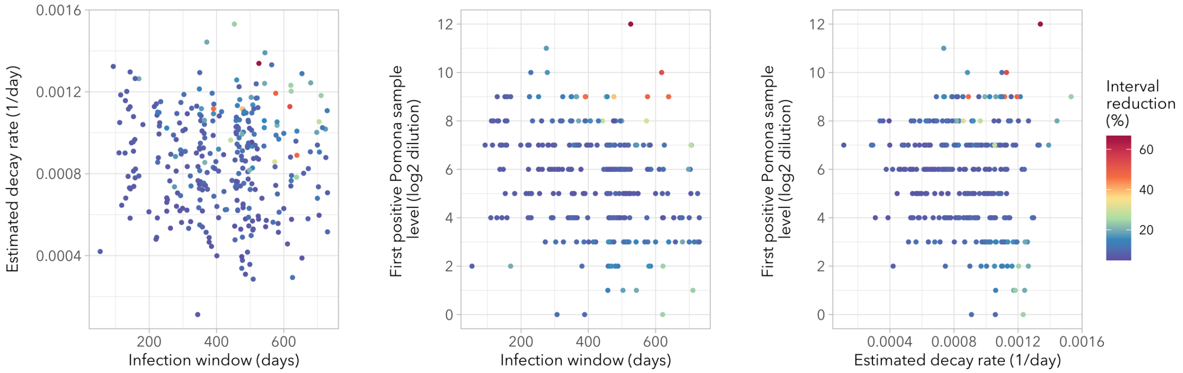


Figure S9. **Infection window reduction (heatmap colors) for key correlates decay rate, infection window size and level of the first positive sample**.

### 4. Effect of peak antibody level and decay rate on infection window size

Table S5. **Effect of peak antibody level and decay rate on infection window estimation**. Gained information on the size of the infection window relative to the prior knowledge, which is a uniform interval bound by the last negative and first positive samples. Left: the percentage by which the infection window was reduced, where the 95% CrI of the posterior distribution was taken as the new window. Right: the information gained by the posterior distribution, expressed as relative entropy, where a higher value indicates a larger information gain expressed in 'bits' units.

| Infection window reduction (%) | | | | | Relative entropy (bits) | | | | |
|---|---|---|---|---|---|---|---|---|---|
| **Peak Ab SD ($log_2$ units)** | **Decay rate ($log_2$ units/day)** | | | | **Peak Ab SD ($log_2$ units)** | **Decay rate ($log_2$ units/day)** | | | |
| | **0.005** | **0.001** | **0.0005** | **0.0001** | | **0.005** | **0.001** | **0.0005** | **0.0001** |
| **0.5** | 49 | 52 | 39 | 10 | **0.5** | 1.04 | 1.07 | 0.71 | 0.11 |
| **1.0** | 48 | 40 | 27 | 8 | **1.0** | 1.01 | 0.75 | 0.44 | 0.07 |
| **2.0** | 41 | 30 | 17 | 7 | **2.0** | 0.75 | 0.51 | 0.23 | 0.05 |
| **3.0** | 37 | 21 | 13 | 6 | **3.0** | 0.66 | 0.33 | 0.17 | 0.04 |

### 5. Correlations between peak antibody level and covariates

We tested whether observed peak antibody level correlated with certain variables, as such a correlation could be integrated into a model of peak antibody time and improve the posterior estimate. Peak antibody level was approximated by the level of the first positive sample for the subset of 54 individuals that had been infected recently (described above). Variables tested were: sex, body weight, body condition, age class, reproductive status, birth year and sample collection month. Correlations were tested for each variable using a linear model with normal error distribution, using $log_2$ antibody level as outcome variable. The linear models were fitted using rjags (Plummer, 2019), with uninformative normal prior $\mathcal{N}(0,2)$ for the effect estimates, which were scaled to mean 0 and standard deviation 1 when numeric.

There were no variables for which the 95% credible intervals did not include 0.

The same analysis was performed using peak antibody level estimates from the models instead of observed values for recently negative individuals. Similarly, no statistically meaningful effects were observed.

### 6. Peak antibody levels of serovars Pomona and Autumnalis

Because a preliminary analysis of the antibody levels for serovars Pomona and Autumnalis suggested a linear correlation, we explicitly incorporated this into the Bayesian model (see below) to test whether this was indeed the case. This was done using a multivariate normal distribution for the peak antibody levels of both serovars. The model indeed found a strong correlation between the two (Figure S9). where peak antibody levels for serovar Pomona are on average 0.67 $log_2$ units lower than those for serovar Autumnalis, with a slope of 0.99 (95% CrI 0.90 to 1.14) and an $R^2$ value of 0.99 (95% CrI 0.94 to 1.00).

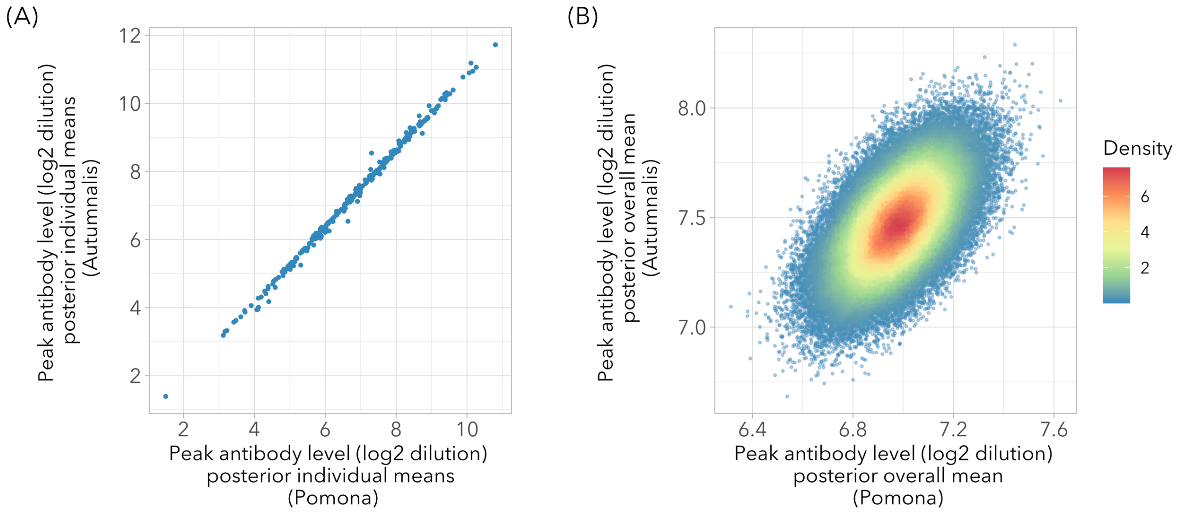


Figure S10. **Correlation between peak antibody levels for serovars Pomona and Autumnalis**. (A): Posterior means of each individual. (B) Posterior MCMC samples of the overall means of peak antibody level for the last 20,000 iterations.

### 7. Estimating time of infection for individuals lacking prior negative samples, and from cross-sectional data

We used the model, with parameters estimated from the main data set, to estimate time of infection for individual foxes that lacked the serological data needed for inclusion in the main study. Posteriors for peak antibody time are shown for individuals that did not have a negative sample preceding their first positive (Figure S11), and for individuals that had only one positive sample (Figure S12). In both cases a bound of 6 years (2190 days) was given for the infection window when a negative sample preceding the first positive was absent. A time of 6 years was chosen because it is longer than any observed infection window, but not so long that it spans an unrealistic time period are a mix of age classes. The situation where only a single positive sample is used resembles that of cross-sectional data, illustrating that the method can be applied to this common situation after training the model using a subset for which multiple samples are available.

Last, we tested whether model performance (measured as window size reduction) correlates with the level of the first positive sample and the original window size, when only the first positive sample is used for estimating time of infection (using only individuals with a negative preceding sample). We found no meaningful correlation, mainly due to the fact that original window size and the level of the first sample were correlated, resulting in an interaction that is not possible to disentangle (Figure S13).

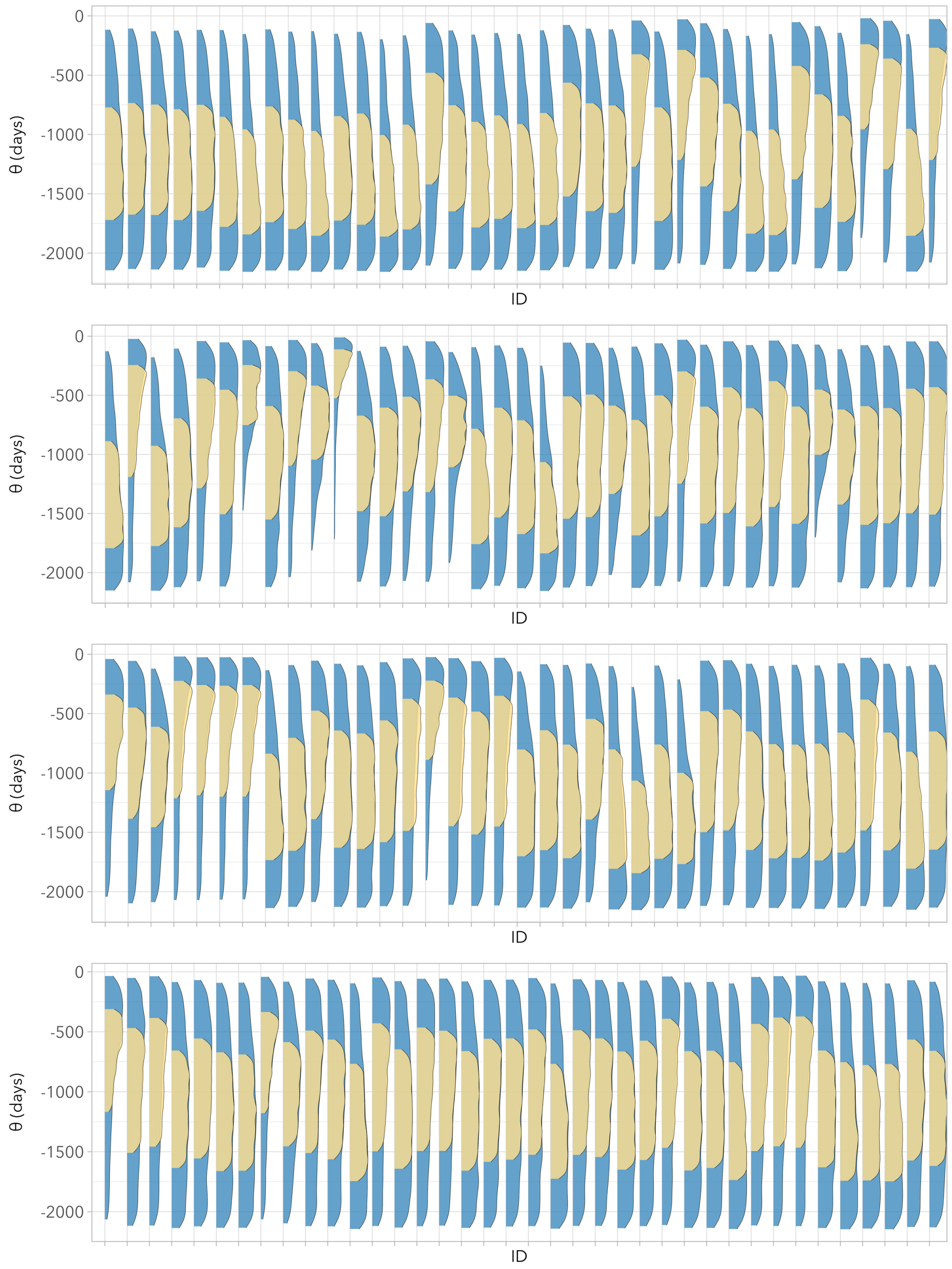


Figure S11. **Infection window posterior densities for individuals without a negative sample preceding the first positive sample.** Colors indicate the size of the credible intervals (blue = 95%, yellow = 50%). Arbitrarily chosen infection windows up to 2190 days (6 years) were possible.

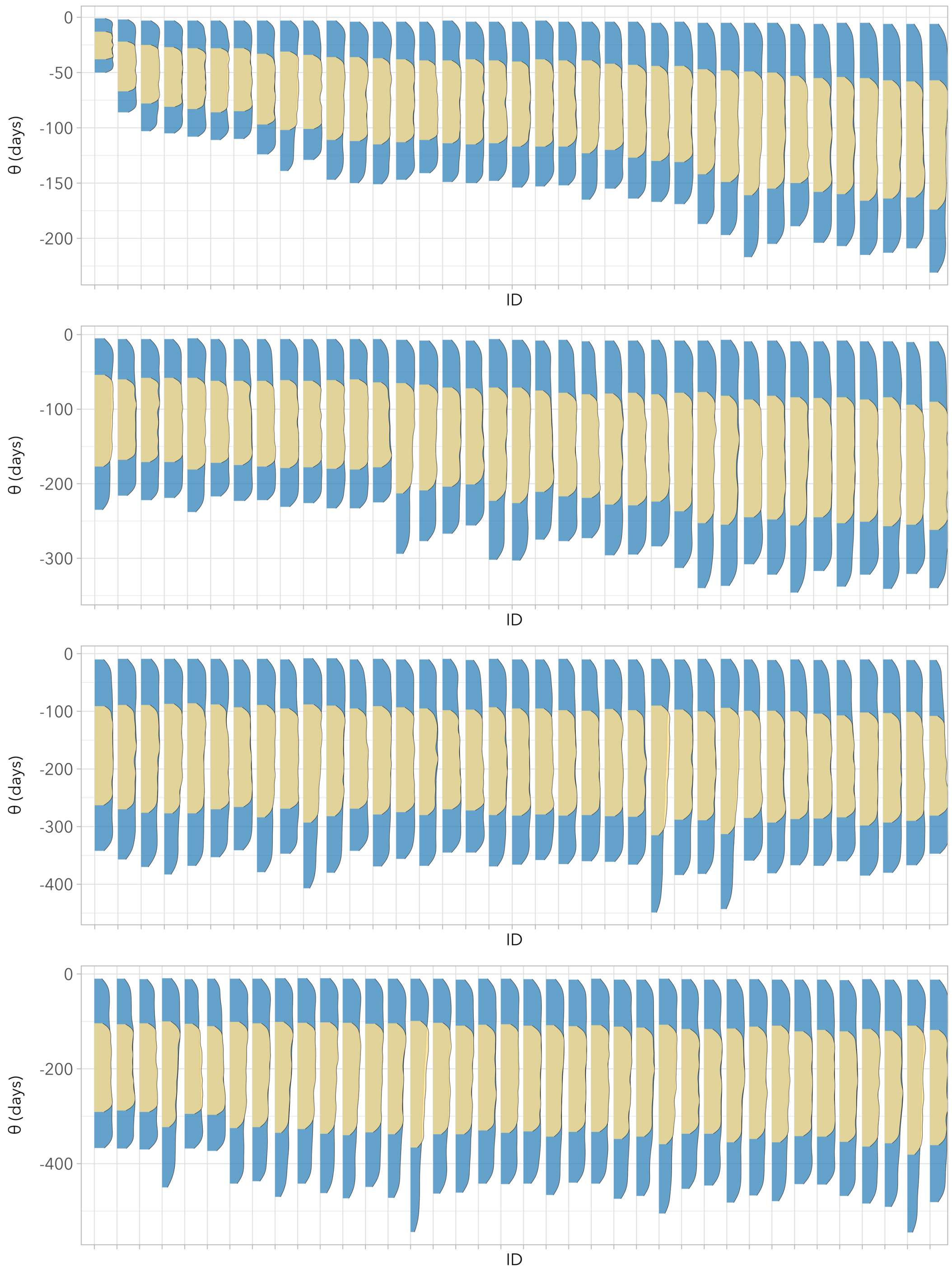


Figure S12. **Infection window posterior densities for individuals with only a single positive sample.** Colors indicate the size of the credible intervals (blue = 95%, yellow = 50%). Arbitrarily chosen infection windows up to 2190 days (6 years) were possible. Only individuals with a preceding negative sample are shown (others are included in Figure S11). Densities are sorted by posterior median.

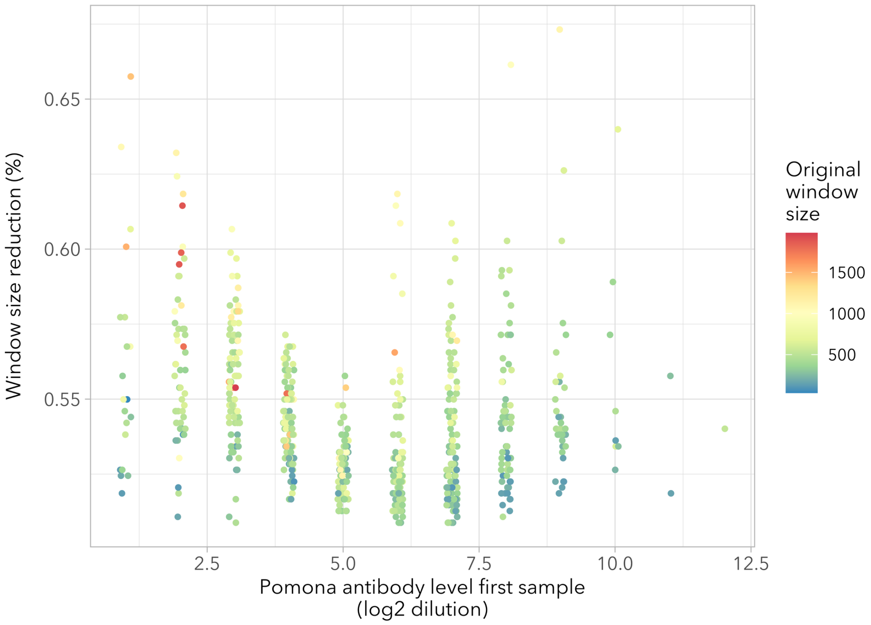


Figure S13. **Infection window size reduction using only the first positive sample for each individual.** Reduction in infection window size plotted against Pomona antibody level ($log_2$ dilution), with colors indicating the size of the original infection window size (i.e. time between the first positive sample and the last preceding negative).

### 8. JAGS code

JAGS code for the double exponential model. A fully working version of this code is also provided as an R markdown document.

```
# priors
for(j in 1:length(neg.int)){

# multivariate distribution for pomona and autumnalis peak antibody levels
mu_pom_aut[j,1:2] ~ dmnorm(mu_pom_aut_mean,mu_pom_aut_precision)

# extract mean peak levels for pomona and autumnalis
mu_pomona[j] <- mu_pom_aut[j,1]
mu_aut[j] <- mu_pom_aut[j,2]

# decay rates pomona and autumnalis
decay_pomona[j] ~ dnorm(decay_overall_pomona,decay_tau_overall_pomona)
decay_aut[j] ~ dnorm(decay_overall_aut,decay_tau_overall_aut)

# time between peak level and first positive, shared between pomona and autumnalis
theta[j] ~ dunif(neg.int[j],0)
}

sigma_pomona ~ dunif(0,50)
tau_pomona <- 1/(sigma_pomona*sigma_pomona)

sigma_aut ~ dunif(0,50)
tau_aut <- 1/(sigma_aut*sigma_aut)

lab_effect ~ dnorm(0,0.01)


#hyper priors

# multivariate pomona autumnalis mean and sd
mu_pom_aut_mean ~ dmnorm(mu_means,tau_means)
mu_pom_aut_precision ~ dwish(omega,wishdf)

# extract individual means peak level
mu_overall_pomona <- mu_pom_aut_mean[1]
mu_overall_aut <- mu_pom_aut_mean[2]

# multivariate precision matrix
inverse_mu_pom_aut_precision <- inverse(mu_pom_aut_precision)
sigma_overall_pomona <- inverse_mu_pom_aut_precision[1,1]^(1/2)
sigma_overall_aut <- inverse_mu_pom_aut_precision[2,2]^(1/2)

# decay rates
decay_overall_pomona ~
dgamma(decay.rate.mean.prior.shape.pomona,decay.rate.mean.prior.rate.pomona)
decay_sigma_overall_pomona ~
dgamma(decay.rate.sd.prior.shape.pomona,decay.rate.sd.prior.rate.pomona)
decay_tau_overall_pomona <- 1/(decay_sigma_overall_pomona*decay_sigma_overall_pomona)

decay_overall_aut ~ dgamma(decay.rate.mean.prior.shape.aut,decay.rate.mean.prior.rate.aut)
decay_sigma_overall_aut ~ dgamma(decay.rate.sd.prior.shape.aut,decay.rate.sd.prior.rate.aut)
decay_tau_overall_aut <- 1/(decay_sigma_overall_aut*decay_sigma_overall_aut)
```

```
# likelihood
for(i in 1:length(time)){
# predicted level pomona
titer_pred_pomona[i] <- lab_effect*lab[i] + mu_pomona[id[i]]*exp(-
decay_pomona[id[i]]*(time[i]-theta[id[i]]))
true_titer_pomona[i] ~ dnorm(titer_pred_pomona[i],tau_pomona)

# interval censoring
titer_pomona[i] ~ dinterval(true_titer_pomona[i], c(1,2,3,4,5,6,7,8,9,10,11,12,13))

# predicted level autumnalis
titer_pred_aut[i] <- lab_effect*lab[i] + mu_aut[id[i]]*exp(-decay_aut[id[i]]*(time[i]-
theta[id[i]]))
true_titer_aut[i] ~ dnorm(titer_pred_aut[i],tau_aut)

# interval censoring
titer_aut[i] ~ dinterval(true_titer_aut[i], c(1,2,3,4,5,6,7,8,9,10,11,12,13))

# store sum loglikelihood as parameter for WAIC calculation
LogLik[i] = log(dnorm(titer_pomona[i], true_titer_pomona[i],tau_pomona)) +
log(dnorm(titer_aut[i], true_titer_aut[i],tau_aut))
}
```
